# Shadow ORFs illuminated: long overlapping genes in *Pseudomonas aeruginosa* are translated and under purifying selection

**DOI:** 10.1101/2021.02.09.430400

**Authors:** Michaela Kreitmeier, Zachary Ardern, Miriam Abele, Christina Ludwig, Siegfried Scherer, Klaus Neuhaus

**Affiliations:** Chair for Microbial Ecology, TUM School of Life Sciences, Technische Universität München Weihenstephaner Berg 3, 85354 Freising, Germany; Bavarian Center for Biomolecular Mass Spectrometry (BayBioMS), TUM School of Life Sciences, Technische Universität München, Gregor-Mendel-Strasse 4, 85354 Freising, Germany; Core Facility Microbiome, ZIEL – Institute for Food & Health, Technische Universität München Weihenstephaner Berg 3, 85354 Freising, Germany

**Keywords:** Overlapping genes, mass spectrometry, ribosome profiling, orphan genes, overprinting, *de novo* gene origin

## Abstract

The existence of overlapping genes (OLGs) with significant coding overlaps revolutionises our understanding of genomic complexity. We report two exceptionally long (957 nt and 1536 nt), evolutionarily novel, translated antisense open reading frames (ORFs) embedded within annotated genes in the medically important Gram-negative bacterium *Pseudomonas aeruginosa*. Both OLG pairs show sequence features consistent with being genes and transcriptional signals in RNA sequencing data. Translation of both OLGs was confirmed by ribosome profiling and mass spectrometry. Quantitative proteomics of samples taken during different phases of growth revealed regulation of protein abundances, implying biological functionality. Both OLGs are taxonomically highly restricted, and likely arose by overprinting within the genus. Evidence for purifying selection further supports functionality. The OLGs reported here are the longest yet proposed in prokaryotes and are among the best attested in terms of translation and evolutionary constraint. These results highlight a potentially large unexplored dimension of prokaryotic genomes.

## Introduction

The tri-nucleotide character of the genetic code enables six reading frames in a double-stranded nucleotide sequence. Protein-coding ORFs at the same locus but in different reading frames are referred to as overlapping genes (OLGs). Studies of coding overlaps of more than 90 nucleotides, i.e. non-trivial overlaps, have mainly been restricted to viruses, where the first OLG was found in 1976^1^. An OLG pair with such a non-trivial overlap can be described in terms of an older, typically longer “mother gene” and more recently evolved “daughter” gene by analogy to mother and daughter cells in reproduction.

In prokaryotes, automated genome-annotation algorithms like Glimmer allow only one open reading frame (ORF) per locus with the exception of only short overlaps^2^ This systematically excludes overlapping ORFs from being annotated as genes^3^. The ‘inferior’ ORFs (e.g. shorter or fewer hits in databases) within overlapping gene pairs have been called ‘shadow ORFs’ since they are found in the shadow of the annotated coding ORF^4^. Determining which ORF to annotate within such pairs has been described as the most difficult problem in prokaryotic gene annotation^5^. Nevertheless, a few prokaryotic OLGs have been discovered, often serendipitously in the pursuit of other unannotated genes^6–8^. For instance in some *Escherichia coli* strains a few non-trivial overlaps have been detected and experimentally analysed^9–16^. Transcriptomic or translatomic evidence for OLGs also exist in genera such as *Mycobacterium*^17^ or *Pseudomonas*^18^. Recently, antisense OLGs have also been reported in archaea^19^ and in mammals^20,21^. These findings support the hypothesis that OLGs are ubiquitous.

Verifying OLGs using mass spectrometry (MS) is difficult because most OLGs appear to be short and weakly expressed, and proteomics has limited abilities in detecting such proteins^22^. RNA sequencing led to the discovery of many antisense transcripts, but whether many of these are translated is controversial^23^. More recently, mRNA protected by ribosomes after enzymatic degradation has been sequenced using “ribosome profiling”^24^ (RiboSeq), showing evidence of translation for antisense transcripts^16,23^. However, artefacts may occur due to structured RNAs^25^. Moreover, proteins produced from such transcripts have been claimed to be predominantly non-functional in *Mycobacterium tuberculosis*^17^. Detecting purifying selection would provide evidence for functionality in OLGs; however, this is made difficult by selection pressure on the other reading frame. Methods have recently been developed^26–30^, but so far applied mainly in viruses. One exception is a cursory study of some OLG candidates in *E. coli*^31^. Additional reason to expect to find OLGs under selection in bacteria comes from a study showing excess long antisense ORFs in a pathogenic *E. coli* strain^32^. Another is the frequent exchange of functional genes between phages and bacteria^33^.

The evolution and origin of OLGs and their constraints have long been discussed^34–37^ and were briefly mentioned in two early books^38,39^. The origin of a new gene within an existing gene is termed “overprinting”^40^. Advantageous for same-strand overlaps is that the hydrophobicity profile of a frame-shifted sequence tends to be similar to that of the unshifted sequence^41^, as may have contributed to the origin of an OLG in SARS-CoV-2^42^. Intriguingly, overlaps of the reference frame “+1” and antisense “−1” tend to have opposite hydrophobicities for their amino acids. Further, similar amino acids are conserved in the antisense frame following synonymous mutation in the reference frame^43^, facilitating the maintenance of overlapping genes. The developing research area of overlapping gene origins complements recent findings of many taxonomically restricted (“orphan”) genes^44^ and unannotated short genes in prokaryotes^45^.

The genus *Pseudomonas* (Gram-negative, Gammaproteobacteria) is of particular interest regarding long OLGs. Its high GC content of 60-70% results in longer average lengths for antisense ORFs compared to *Escherichia* (~50% GC)^32^. Previous studies reported OLGs in the well-characterized species *P. fluorescens* and *P. putida*, although based on limited evidence^46–48^. Since then, methods allowing improved discovery and verification of such genes have been developed. Here, we examine *Pseudomonas aeruginosa*, which is a versatile pathogenic species^49^. As an opportunistic human pathogen, it predominately causes disease in immunocompromised individuals^50^. Hospital-acquired infections with *P. aeruginosa* are often associated with high morbidity and mortality and may include infections of the respiratory tract, the urinary tract, the bloodstream, the skin, and wounds^51^. The high level of intrinsic and acquired resistance as well as the rapid rise of multi-drug resistant strains limit effective antibiotic treatment options^52,53^. Thus, it is important to understand the entire genomic complexity of this organism. In *P. aeruginosa,* we detected two exceptionally long novel ORFs showing large overlaps of 957 nt and 1542 nt in antisense to annotated genes, by employing RiboSeq as well as mass spectrometry. We present evidence that both OLGs indeed encode proteins which evolved only recently by overprinting. Further, we detected purifying selection operating through a depletion of stop codons and non-synonymous changes.

## Results

### Genomic locations of two long overlapping genes

The novel *olg1* of *P. aeruginosa* PAO1 (NC_002516.2) is located at the coordinates 291556-292512(+) in frame −1 (i.e. directly antisense) with respect to its mother gene (Fig. 1a). It has a minimum length of 957 nucleotides (nt) and the most probable start codon is ATG_291556_ (see Supplementary Information). *olg1* completely overlaps with the annotated mother gene *tle3* (PA0260), encoding the toxic type VI lipase effector 3 of the *vgrG2b-tli3-tle3-tla3* operon. *Tle3* contains two structural domains, an N-terminal α/β hydrolase fold domain and an C-terminal domain of unknown function (DUF3274)^54^. *Olg1* overlaps 39 nt with the α/β fold domain and at least 594 nt with DUF3274.

**Fig. 1.**
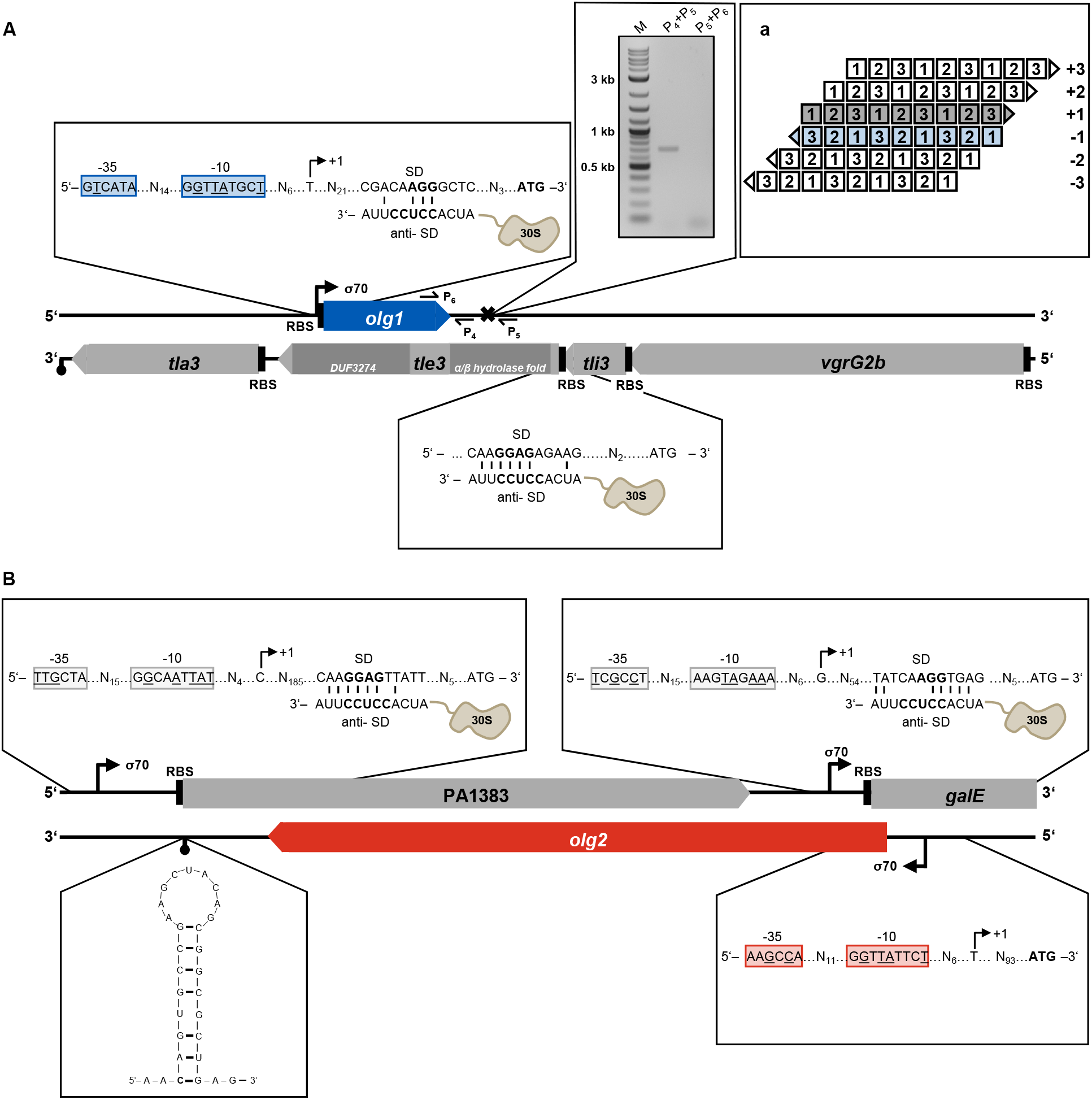
Schematic overview of the genomic structure of the *tle3*-*olg1* (A) and PA1383 *-olg2* (B) locus. **(A)** *olg1* completely overlaps antisense in frame −1 (**a**) relative to the annotated gene *tle3*, which is part of the *vgrG2b-tli3-tle3-tla3* operon. Location of the N-terminal α/β hydrolase fold as well as the C-terminal DUF3274 domain of *tle3* are displayed in dark grey. *olg1* shares structural features of a protein-coding gene including −1 and −1 consensus elements, divided by a 14 bp spacer, of a putative σ^70^ promoter. A core SD sequence of AGG was identified according to Ma et al. ^61^, interacting with the aSD sequence at the 3’ end of the 16S rRNA within the 30S ribosomal subunit. A putative terminator between 219 and 349 nt downstream of the stop codon was identified via RT-PCR using the primer pairs indicated (P_4/5_+P_6_). **(B)** *olg2* overlaps non-trivially with the hypothetical gene PA1383 and trivially with *galE* encoding a UDP-glucose 4-epimerase. Structural features of both annotated genes are indicated. The mRNA of *olg2* starts probably at a putative σ^70^ promoter 93 nt upstream of the start codon and terminates at the predicted terminator 218 to 247 nt downstream of the stop codon.

Novel *olg2* (Fig. 1b) is likewise encoded in frame −1, in the mother gene PA1383 at the coordinates 1501875-1503602(−). With ATG_1503602_ as putative start codon, *olg2* spans 1728 nt and overlaps with two annotated genes. A region of 1536 nt is shared with PA1383, a hypothetical gene predicted to code for an N-terminal Type I signal sequence for cytoplasmic export^55^, which has been shown to be regulated by the transcriptional repressor NrdR^56^, and by both the repressors mvaT and mvaU^57^. Additionally, *olg2* overlaps in frame +2 by 34 nt with the *galE* gene (PA1384), which encodes for a UDP-glucose 4-epimerase.

Both OLGs were discovered by screening RiboSeq data and further characterized using prediction tools, quantitative PCR, transcriptome sequencing, mass spectrometry as well as phylogenetic analyses as described in the following sections.

### *Ab-initio* prediction indicates gene-like sequence features

Putative σ^70^ promoter sequences were searched for 300 nt upstream of each OLG’s start codon with the tool BROM^58^. Linear-discriminant function values of 1.94 (*olg1*) and 1.37 (*olg2*) clearly exceed 0.2, the threshold distinguishing promoter and non-promoter sequences. Transcription start sites were localized 37 nt and 94 nt upstream of the start codons, respectively (Fig. 1). The observed distances fit the length of 5’UTRs reported for *P. aeruginosa* PA14 (median: 47 nt)^59^ and *P. syringae* pv. *tomato* str. DC3000 (mean: 78 nt)^60^. To investigate potential ρ-independent terminators, FindTerm^58^ was applied 300 nt downstream of both overlapping ORFs. A terminator was detected for *olg2* 218 to 247 nt downstream (Fig. 1b). For *olg1* a ρ-independent terminator was not predicted, but RT-PCR verified termination between 219 and 349 nt downstream of its stop codon.

In *P. aeruginosa*, the core anti-Shine-Dalgarno (SD) sequence is CCUCC. It has a mean *ΔG*_*SD*_ of −6.5 kcal/mol and an optimal spacing of 7-9 nt to the start codon^61^. A SD sequence with AGG and a *ΔG*_*SD*_ of −3.6 kcal/mol was detected 8 nt upstream of *olg1*′s proposed start codon (Fig. 1a). For the mother gene *tle3*, a SD sequence (−5.1 kcal/mol) was identified, but neither a σ^70^ promoter nor a ρ-independent terminator was found. This is consistent with the reported finding that *tle3* is part of the *vgrG2b*-operon^54^. No SD sequence was detected for *olg2*. However, SD sequences are similarly absent from 30.8% of the annotated genes in *P. aeruginosa*^61^. The upstream region of both PA1383 and *galE*, the mother genes, harboured a σ^70^ promoter (transcription start sites 203 nt and 72 nt upstream, respectively) and a SD sequence (−6.1 kcal/mol and −4 kcal/mol, respectively).

Sequence features such as start codon, SD sequences, GC bias, and hexamer coding statistics are used by gene prediction tools, for example Prodigal^62^. However, Prodigal’s algorithm prohibits prediction of gene pairs with an overlap larger than 200 nt and annotates only the ORF with the highest score. For *P. aeruginosa* PAO1, Prodigal predicted 5681 protein coding genes including the two annotated genes *tle3* and PA1383 with total scores of 304.76 and 225.92, respectively (Supplementary Fig. 1). When “hiding” both annotated genes by replacing all start codons by unidentified nucleotides (“N”), Prodigal classified *olg1* and *olg2* as protein coding genes. While their total scores of 4.63 and 23.63 were relatively low, some values are comparable to annotated genes, for instance the start-sequence region scores (Supplementary Fig. 1 and Supplementary Table 1). Furthermore, *olg2* showed a confidence score of 99.56, indicating a very high likelihood of being a real protein-coding gene. These overlapping ORFs are not annotated by Prodigal due to their long overlaps but both nonetheless show features associated with protein coding.

### RNASeq and RiboSeq confirm gene expression

Transcription and translation of the overlapping ORFs and their mother genes was analysed by RNASeq and RiboSeq^24^. *P. aeruginosa* PAO1 was cultivated under standard conditions (LB, 37°C) in two biological replicates. RNASeq revealed high RPKM_RNASeq_ values between 20 (PA1383) and 36 (*olg1*) for all four ORFs (Fig. 2ab, first track; Supplementary Table 2). These values were comparable to annotated protein-coding genes which range between 0.4 and 13563.5 (Supplementary Fig. 2a), indicating substantial transcription. Full-length transcription of both OLGs was confirmed by RT-PCR (Supplementary Fig. 3). Antisense transcription, reported in diverse bacteria including *P. aeruginosa*, often plays a role in gene regulation, indicated by a negative correlation between antisense and mRNA levels^63,64^. However, evidence exists that some antisense transcripts are translated^8,23,65^. Our results (Fig. 2ab, second track) strongly support translation in both directions for the loci of PA1383 and *tle3*. The annotated genes showed RPKM_RiboSeq_ values close to the median annotated gene RPKM_RiboSeq_ of 25.7 (Supplementary Fig. 2b). The expression of *olg1* was even higher (RPKM_RiboSeq_ = 40.3), whereas *olg2* showed a lower, but still unequivocal signal (RPKM_RiboSeq_ = 14.2). In each case expression was consistent across replicates and coverage was excellent across the whole ORF (Supplementary Fig. 2c and Fig. 2ab, second track). The structural features reported above overall matched the read distributions. However, reads upstream of the proposed *olg1* start codon ATG_291556_ suggest additional transcription and translation (question mark in Fig. 2a), perhaps from an alternative start codon or a different frame ORF upstream (Fig. 2a, first and second track).

**Fig. 2.**
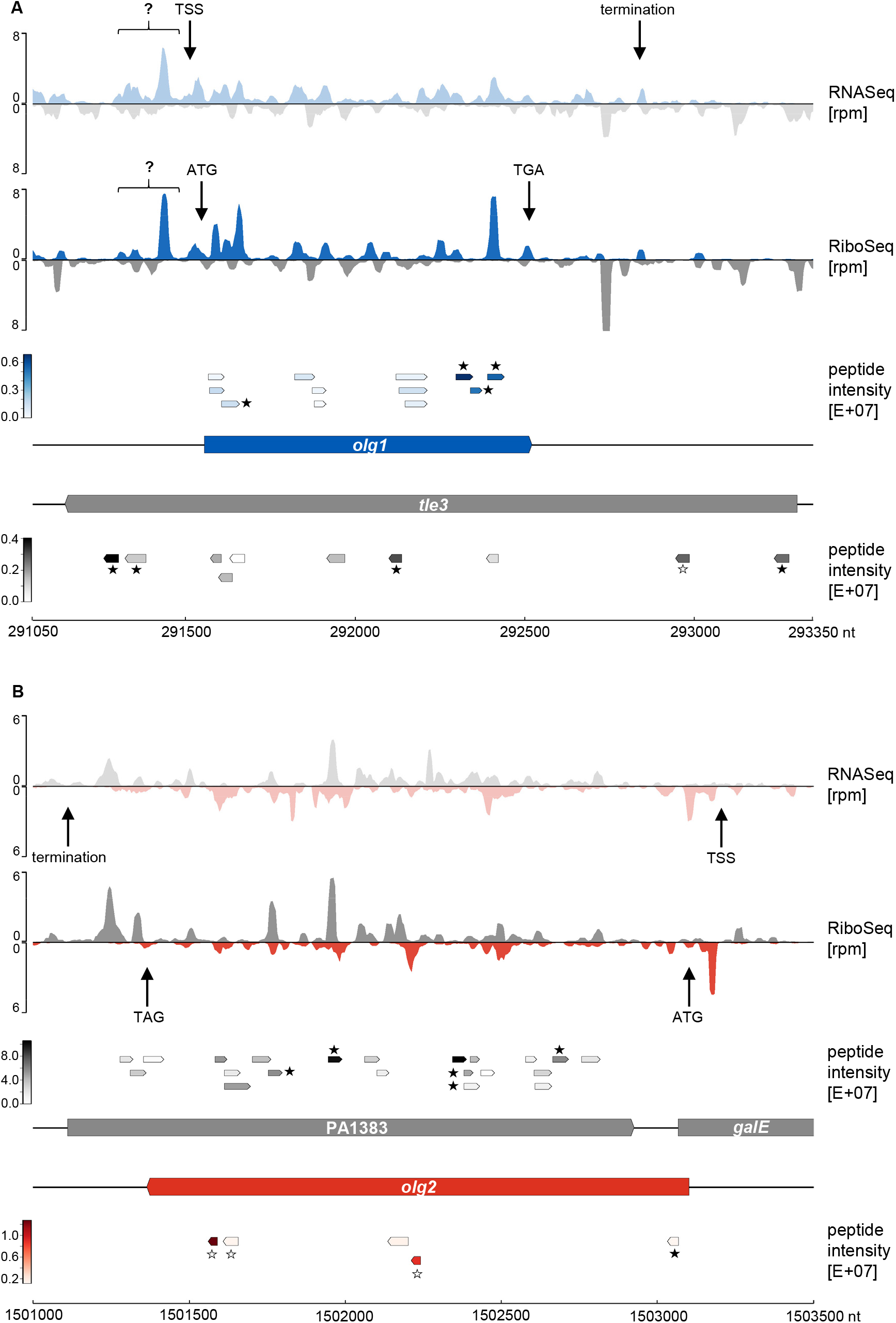
RNASeq, RiboSeq and mass spectrometry signals at the *olg1*-*tle3* (A) and *olg2*-PA1383 (B) locus. Shown are the mean normalized rpm values of all transcriptome (first track) and translatome reads (second track) of this study (n=2) for *olg1* (blue), *olg2* (red) and their mother genes *tle3* and PA1383 (both grey). Transcription start (TSS) and stop sites (termination) as well as the positions of start and stop codons are indicated by arrows. Signals of unknown origin upstream of *olg1* are highlighted by a question mark. Track 3 and 4 illustrate the position and intensity of all peptides obtained by mass spectrometry. Peptides that were selected for targeted proteomics (PRM) are highlighted with an asterisk. Peptides validated and quantified by PRM are indicated by filled asterisks

DeepRibo is a software combining RiboSeq signals and DNA sequence motifs by using neural networks^66^. DeepRibo predicts *olg1* and *olg2* to be translated with the same start codons as we predict from visual assessment, in both replicates; it also confirms translation of the mother genes *tle3* and PA1383. As a comparison, we analysed the genes adjacent to *tle3* - these genes similarly had high coverage and RPKM values (Supplementary Table 2), implying expression of the operon, but no antisense signals (Supplementary Fig. 4). Finally, ribosome coverage values (RCV) were calculated, i.e. RPKM_RiboSeq_ over RPKM_RNASeq_^67^. This value allows a direct estimation of the “translatability” of an ORF. With a value of 1.13, *olg1* and PA1383 were within the top 20% of all annotated genes. *tle3* and *olg2* showed lower RCVs (i.e., 0.79 and 0.64, respectively) but within the range of other annotated genes (Supplementary Fig. 2d).

### Mass spectrometry identifies translated peptides

Mass spectrometry-based proteomics was used to verify the expression of both OLGs as well as the respective mother proteins. For that, *P. aeruginosa* PAO1 was cultivated as described above. In total 4,076 proteins could be detected, including Olg1, Olg2, Tle3 and PA1383 with 12, 5, 10, and 21 peptides, respectively (Supplementary Table 3). These peptides covered the start, middle and end regions in each of the four coding sequences (Fig. 2, last tracks). The measured mass spectrometric intensities of the four target proteins were in a medium to low range compared to all other detected proteins (Supplementary Fig. 5). To exclude false-positive peptide identifications, we validated the fragment ion spectra of all detected peptides from the four target proteins using the artificial intelligence algorithm Prosit^68^. Prosit can predict a peptide’s fragment ion spectrum based on its amino acid sequence. Except for two peptides, correlation scores (dot product) between experimental and predicted spectra were larger than 0.6 (Supplementary Table 3), which supports correct identification of almost all putative peptides from Olg1, Olg2, Tle3 and PA1383.

For peptide identification with highest confidence, as well as for accurate protein quantification, we next performed targeted proteomic measurements including isotopically labelled reference peptides. Based on our initial mass spectrometric data, we selected four to five peptides per target protein (Fig. 2, last tracks, peptides indicated with an asterisk) and purchased those in synthetic and stable isotopically-labelled form. Those heavy reference peptides were spiked into *P. aeruginosa* PAO1 samples taken from various growth time points (Fig. 3a) and measured using the targeted proteomic method Parallel Reaction Monitoring (PRM). We successfully validated and quantified 4 peptides for Olg1, 5 peptides for PA1383, 4 peptides for Tle3, and one peptide of Olg2 (Fig. 3b and Supplementary Fig. 6a).

**Fig. 3.**
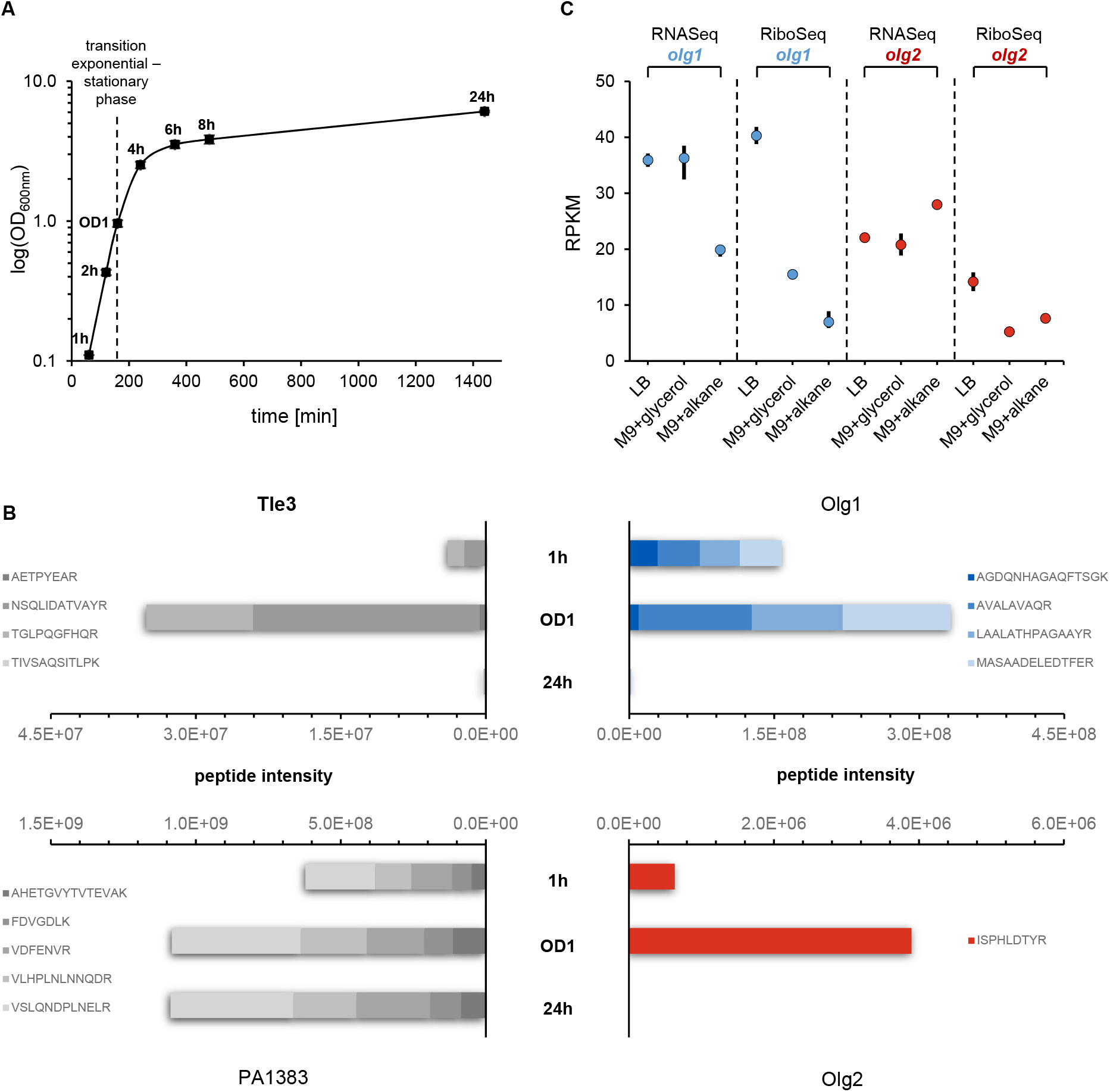
Regulated protein expression of Olg1 and Olg2. **(A)** Shownare the mean OD_600nm_ values measured for *P. aeruginosa* PAO1 in three biological replicates. Samples were taken for targeted proteomics and qPCR at 1h, 2h, 4h, 6h, 8h and 24 h as well as at OD_600nm_=1 (~160 min) as indicated. **(B)** Peptideintensities measured by targeted proteomics (PRM) for the proteins Tle3, Olg1, PA1383 and Olg2 at selected time points. **(C)** Meantranscriptome and translatome reads per kilobase per million mapped reads (RPKM) of the datasets “LB” (n=2; this study) as well as “M9+glycerol” and “M9+alkane” (n=3 each; published by Grady et al.^69^) are shown for *olg1* (blue) and *olg2* (red).

### Regulation of gene expression implies functionality

Growth-phase dependent changes in protein abundances were observed for all four target proteins via PRM (Fig. 3b and Supplementary Fig. 6a). High levels of both OLGs were obtained during exponential growth (1 h, 2 h) and at the exponential-stationary transition (OD1). In contrast, protein abundance in late stationary phase (24 h) was significantly lower. Both OLGs exhibited protein kinetics deviating from their respective mother gene proteins Tle3 and PA1383, indicating an independent biological regulation. qPCR analysis confirmed similar kinetics for the mRNAs of *olg1* and *olg2* (Supplementary Fig. 6c).

Further indication of regulated OLG-expression was provided by published RNASeq and RiboSeq data in two *P. aeruginosa* strains (PAO1 & ATCC33988)^69^. Strains were cultivated in M9 broth with glycerol or *n*-alkanes. Reanalysis revealed transcriptional and translational signals for both OLGs in strain PAO1 (Fig. 3c and Supplementary Fig. 7) and for *olg1* in strain ATCC33988 (Supplementary Table 2). Interestingly, RPKM_RNASeq_ values were similar for both OLGs when comparing LB with M9+glycerol, but differed in M9+alkane (Fig. 3c). This might suggest a carbon-source-dependent regulation. Further, these data indicate regulated translation of *olg1* (log_FC = 1.15,) and *olg2* (log_FC = −0.55) in PAO1 when comparing M9+alkane with M9+glycerol (false discovery rate, FDR ≤0.05). Thus, both OLGs are expressed more weakly in alkane media than LB (Fig. 3c).

Temporal control and dependence on growth media for OLG expression strongly implies functionality (Fig. 3b and c, Supplementary Fig. 6). However, elucidation of the biological role of these overlapping-encoded proteins requires further experiments.

### Phylogenetic analyses show a relatively recent origin

Almost all detectable homologs of both *olg1* and *olg2* were found within *Pseudomonas* spp. according to BLAST searches (Supplementary Table 4). Homologs outside *Pseudomonas* clustered within the genus in terms of sequence identity. Thus, we infer that these are best accounted for by horizontal gene transfer from the genus and focus on evolution within the genus *Pseudomonas*.

Homologs of *tle3* containing both the α/β hydrolase and DUF3274 domains were found in multiple phyla, however within the order Pseudomonadales they were found only in the genus *Pseudomonas*, suggesting horizontal gene transfer from another order. The mother gene of *olg2*, PA1383, in contrast, does not have highly similar homologs outside of *Pseudomonas*. The one exception was derived from a low-quality genome of *Acinetobacter baumanii* (noted in RefSeq to be of excess size), which was disregarded. A few distant homologs were found, with the two top hits in *E. coli* (matching 43% of the PAO1 sequence) and *Salmonella enterica* (match to 17%). This evolutionary distance again implied a horizontal gene transfer into or out of *Pseudomonas* with subsequent evolutionary divergence. None of the non-*Pseudomonas* homologs included the N-terminal signal peptide in PA1383, suggesting functional changes in *Pseudomonas* compared to other bacteria.

Both *olg1* and *olg2*, in the length present in reference strain PAO1, were highly taxonomically restricted. The taxonomic distribution of both OLGs was limited approximately to the species *P. aeruginosa*, according to BLASTp hits along with additional metagenome assembled genomes (MAGs) (Fig. 4a). The mother genes *tle3* and PA1383 within the order *Pseudomonadales* resulted in approximately 900 and 300 unique sequences, respectively. For *olg1* (in *tle3*), the intact ORF (i.e. without premature stop codons) was restricted to *P. aeruginosa* with one exception in *P. prosekii* with a non-start codon (GTA) at the locus of the start site in PAO1. An intact *olg2* was restricted to a few *Pseudomonas* species, and only *P. aeruginosa* genomes shared the same stop codon. *Pseudomonadales* homologs in recent MAG collections^70,71^ supported the inferred taxonomic boundaries. Intact ORFs of *olg1* or *olg2* with a stop in the same position were assigned taxonomically to *P. aeruginosa* in the MAG data (Supplementary Table 4). Subsequent analyses used the combined GenBank and MAG datasets.

**Fig. 4.**
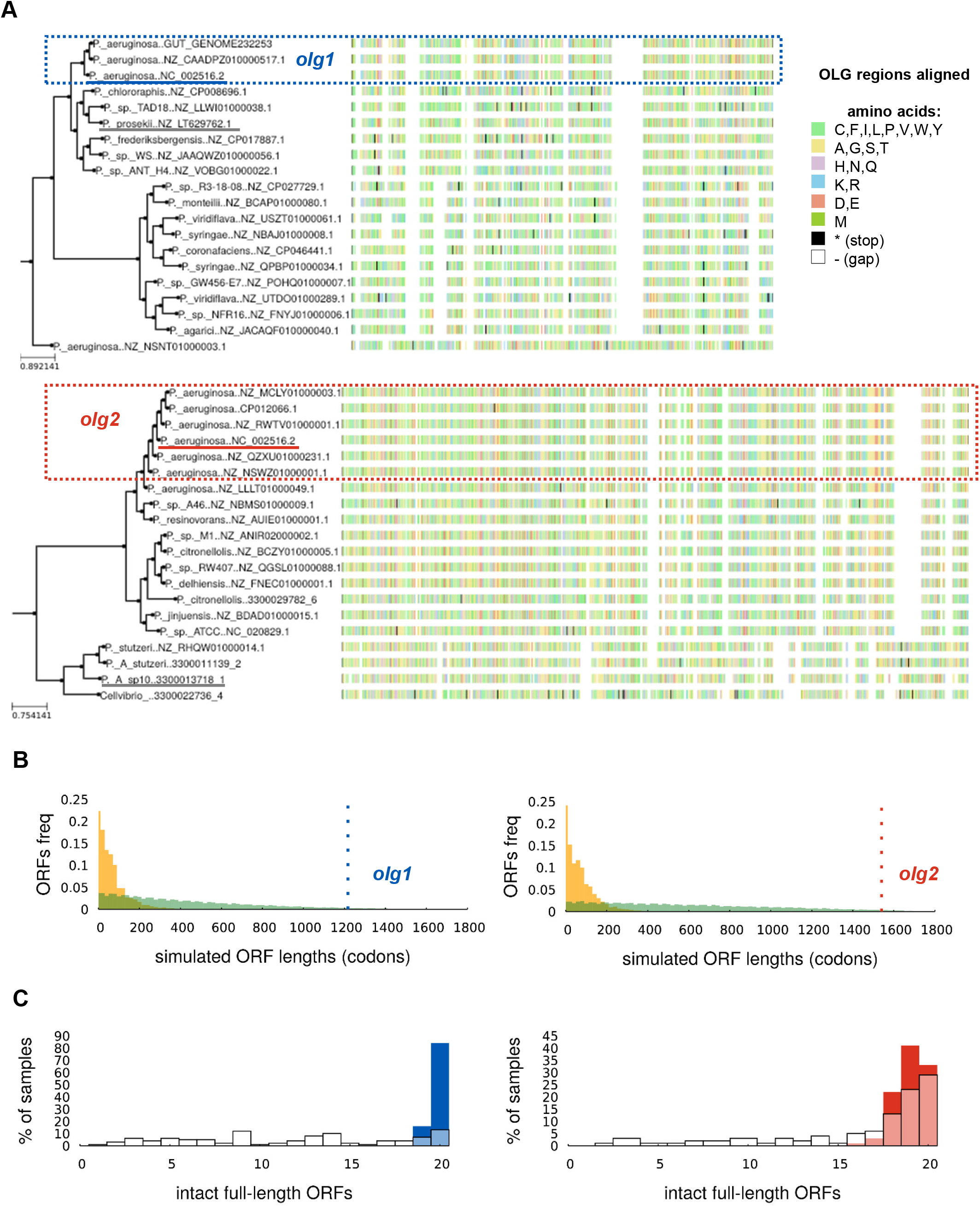
Phylogenetic distribution of *olg1* and *olg2* and depletion of stop codons. **(A)** Homologs of the full *olg1* ORF (left), matched to a maximum likelihood tree calculated from the amino acid sequence of the mother gene *tle3*, down-sampled to 20 genomes. Clade containing genomes with the same start and stop codon as the reference genomes (“OLG genomes”) is highlighted with a blue box. Right: homologs of *olg2* overlapping loci, for PA1383. Clade containing genomes with the same stop codon as the reference genomes is highlighted with a red box (start codon is at a non-overlapping locus outside the sequence shown). The reference genome (NC_002516.2) is underlined in the respective OLG colour, and the outgroup used in the evolutionary simulations described below is underlined in grey. **(B)** Distributions of lengths of antisense (−1 frame) ORFs obtained by permutation (green) or synonymous exchanges (orange) of ‘mother gene’ codons, for genes *tle3* and PA1383, compared to the lengths of the embedded *olg1* (blue, left) and *olg2* (red, right). ORF lengths are measured between in-frame stop codons rather than start to stop. **(C)** Simulations of evolution of *tle3* and PA1383 in the OLG clade rooted on an outgroup with an intact ORF, using an empirical codon model, show that accumulation of stop codons is common; simulated sequences tend to have fewer full-length intact ORFs in the OLG loci and reading frame than real sequences.

### Stop codon depletion implies purifying selection

Multiple independent lines of evidence indicated that both OLGs are under purifying (negative) selection, a strong indicator of functionality, particularly when combined with evidence of expression^72^. Firstly, the ORFs for *olg1* and *olg2* were both significantly longer than expected given the amino acids (AA) in the reference genes *tle3* and PA1383, using the synonymous-mutation method from the tool ‘Frameshift’^73^ (Fig. 4b). This method substitutes synonymous codons randomly and obtains an empirical cumulative distribution function of the resulting ORF lengths in each alternative reading frame. This resulted in a *P*-value of <10^−10^ for both ORFs. *Olg1* and *olg2* were also longer than expected given the overall codon usage (codon-permutation method of ‘Frameshift’), although not statistically significantly, with p = 0.163, and p = 0.0635 respectively (Supplementary Table 5). The *P*-values included a correction for multiple tests, i.e. the number of observed ORFs in this alternate reading frame. The synonymous mutation P-values are still significant after a conservative multiple-tests adjustment of multiplying by the total number of genes, arguably appropriate given the OLGs’ detection with a genome-wide scan. The non-significant results for the codon permutation method are not surprising given that it implicitly depends on stop codons elsewhere in the alternate reading frame, of which there are few due to the length of the OLGs relative to their mother genes. In summary, from these results it can be concluded that the ORF lengths were not simply a result of the overall sequence composition of the mother genes, implying selection for long ORFs via negative selection on stop codons.

Secondly, sequence evolution of each mother gene was modelled without selection for maintaining an overlapping ORF. The presence of stop codons in simulated sequences was then compared to natural sequences. This method was previously used to support an inference to selection on the OLG *asp* in HIV-1^74^. When evolution was simulated using an empirical codon model in Pyvolve^75^ along trees calculated from the mother genes (Fig. 4a), stop codons evolved more frequently in the simulated OLG sequences compared to observed natural sequences. As such, fewer simulated than natural sequences had intact full-length ORFs (Fig. 4c). Following the originators of the method^74^, outgroup sequences without stop codons in the OLG region were chosen to root the tree.

### Codon position-specific constraint supports purifying selection

Synonymous variation in the mother genes *tle3* and PA1383 was reduced over a large part of the OLG region (Fig. 5a, Supplementary Table 6), according to results from ‘FRESCo’^28^. For *tle3*, a comparison of the rates of synonymous evolution in the OLG-containing genomes versus the rest of the alignment using a paired two tailed t-test in non-overlapping adjacent windows of 50 codons over the whole OLG sequence showed increased constraint in the OLG region, with p = 0.086. Results for *olg2* were similar for the last 350 codons of *olg2* (p = 0.03) (Supplementary Table 6), but there was no synonymous constraint towards the end of the mother gene PA1383 from approximately codon 500 onwards. Because the non-overlapping start region of *olg2* is not well conserved across *P. aeruginosa* (Supplementary Table 4), we focused only on the part overlapping PA1383.

**Fig. 5.**
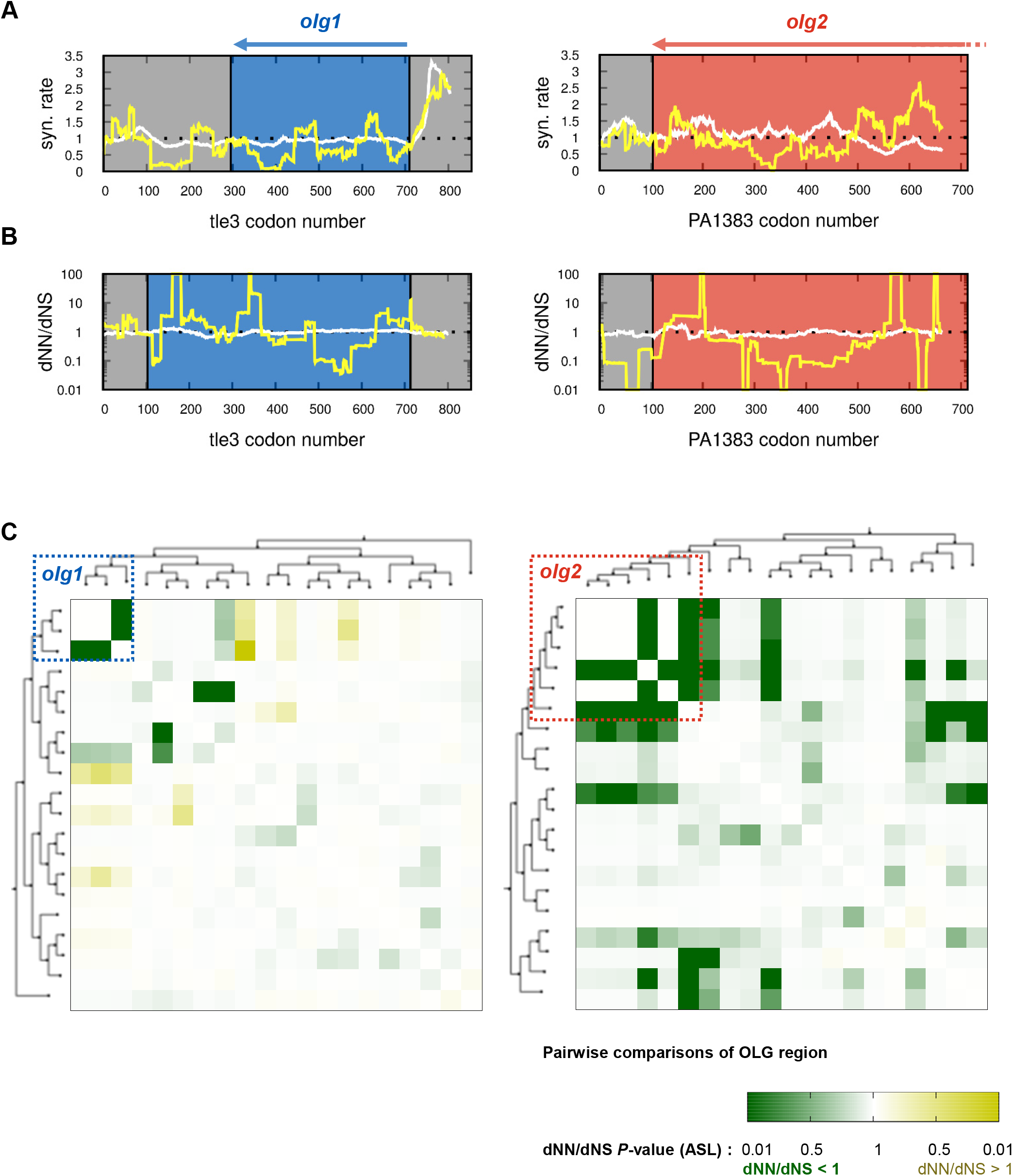
Evidence for evolutionary sequence constraint in *olg1* and *olg2*. **(A)** Variationin synonymous codons of *tle3* (left) and PA1383 (right), sliding windows of 50 codons calculated with FRESCo. Constraint is observed in the OLG regions (blue and red boxes) when compared to the expected rate of 1 (black dotted lines) and to that observed in the “non-OLG” genomes, approximately 1 across the gene (white line). Codon numbers are with respect to an alignment including gaps. **(B)** OLGenie’s measure of dN/dS across *tle3* (left) and PA1383 (right) in the OLG genomes; sliding windows of 50 codons. A decrease in nonsynonymous changes in the OLG frame is observed in the OLG loci (blue and red boxes) when compared to the expected neutral evolution rate of 1 (black dotted lines) and to the non-OLG genomes (white line). **(C)** Pairwise comparisons of dNN/dNS (an OLG-appropriate measure of purifying selection calculated with OLGenie). Evidence for purifying selection is found in a wider taxonomic group than the specific ORFs studied here; for *olg1*, apparent purifying selection is limited to a subclade within *Pseudomonas*, whereas for *olg2* it is found across the genus; for both ORFs however, evidence is strongest in the vicinity of *P. aeruginosa*. Codon numbers are with respect to an alignment including gaps.

A more precise reading-frame-specific measure of purifying selection against non-synonymous variants in OLGs is given by the novel tool ‘OLGenie’^30^. Unlike standard measures of dN/dS, ‘OLGenie’ calculates an OLG-appropriate measure (i.e. dNN/dNS for the OLG) by restricting the analysis to alternative frame sites where variants are non-synonymous in the reference frame. Within the OLG-containing genome sets for both *olg1* and *olg2*, synonymous variants were favoured over variants causing AA changes, although these tendencies were not statistically significant. For *olg1* dNN/dNS = 0.33, p = 0.11, and for *olg2*, dNN/dNS = 0.52, and p = 0.06. These values contrasted with the genomes without *olg1* or *olg2* with dNN/dNS = 1.02, p = 0.66; and dNN/dNS = 0.92, p = 0.20 for *olg1* and *olg2*, respectively (Supplementary Table 7). Both mother genes *tle3* and PA1383 were observed to be under significant purifying selection in the sets of *Pseudomonas* genomes with and without these OLGs. The results from ‘FRESCo’ and ‘OLGenie’ are fully independent, as they depend on synonymous and nonsynonymous sites in the mother genes, respectively. As each measure independently shows a tendency towards constraint, together they provide good evidence of evolutionary constraint in the OLG sequences. Further, some individual pairwise sequence comparisons with ‘OLGenie’ are statistically significant (Fig. 5c, Supplementary Table 7), and these comparisons are informative about the taxonomic extent of functional ORFs. That much of the strongest evidence of purifying selection is taxonomically close to the reference genome PAO1 supports constraint in these OLGs. This was not guaranteed by the presence of an intact ORF in this clade, as any stop codons are excluded from the ‘OLGenie’ analysis; as such, results are not affected by whether an ORF has premature stops. The pattern of purifying selection on *olg1* suggests that a functional ORF may have been found in the common ancestor of *P. aeruginosa* and *P. frederiksbergensis*; the evidence for positive selection between members of this clade and genomes in the other main *Pseudomonas* branch also support this hypothesis. In the case of *olg2*, evidence for purifying selection is taxonomically more widespread across *Pseudomonas*, fitting the wider distribution of intact *olg2* ORFs (some with stop codons downstream of that in the reference strain PAO1).

## Discussion

Prokaryotic OLGs, outside viruses^76,77^, are often categorically rejected and those already annotated have been attributed to misannotations^78^. In this study, we describe the detection and characterization of two exceptionally long OLGs, *olg1* and *olg2*, in *P. aeruginosa*. We propose that the detected overlapping ORFs encode functional protein products due to 1) the presence of structural features necessary for gene expression, 2) successful transcription and translation as indicated by RNASeq and RiboSeq, 3) discovery of several translated peptides via mass spectrometry, 4) validation and confirmation of their regulated expression during growth of *P. aeruginosa* PAO1 using targeted proteomics and isotopically labelled reference peptides, 5) successful prediction of both ORFs on genomic and translational level by annotation programs, and 6) evidence of purifying selection on both gene candidates from multiple methods. While these results provide strong evidence for the genuine protein-coding nature and functionality of both ORFs, they can only be designated as OLGs if their respective mother genes (*tle3* and PA1383) are correctly annotated and are also genuinely protein coding. The gene *tle3* has been confirmed to encode for the antibacterial type VI lipase effector 3^54,79^. PA1383 is annotated as a hypothetical gene, but we show that homologs are widely distributed across bacteria. Further, it contains a signal peptide associated with export, and it is under purifying selection. For both mother genes, we show clear expression in our RNASeq, RiboSeq and MS experiments. It appears unlikely that the MS-detected peptides represent translation products without function considering the high bioenergetic cost of translation^80^. Taken together, it is beyond reasonable doubt that both mother genes encode for functional proteins and that the overlapping ORFs presented here are not just annotation errors from artifactual mother genes.

With a minimum length of 957 and 1728 nt, *olg1* and *olg2* represent the longest known prokaryotic OLGs with extensive experimental evidence. The discovery of such long OLGs is extraordinary considering the short length of most observed OLGs. In *E. coli*, for instance, several RiboSeq studies^8,16,81^ revealed translation of antisense OLGs typically encoding for small proteins, rarely exceeding 200 codons. Similarly, Baek, et al.^7^ detected 130 unannotated ORFs including overlapping ones with total sizes ranging from 4 to 144 AA in *Salmonella enterica* Typhimurium. Likewise, Smith, et al.^17^ report translation of 274 predominately short OLGs in *Mycobacterium tuberculosis*. An equivalent length range was also obtained for antisense OLGs in archaea^19^. The vast majority of OLGs characterized in detail are less than 200 codons^9–14^. Antisense OLGs encoding for proteins equal or larger than 200 AA have rarely been validated experimentally, e.g. *pop* in EHEC (200 AA)^15^, *adm* in *Streptomyces coelicolor* (233 AA)^82^ or *cosA* in *P. fluorescens* Pf0-1 (338 AA)^46^. Even when considering viruses, *olg1* and *olg2* are of exceptional size and belong to some of the longest yet observed OLGs. In a large-scale study of 5,976 viral genomes^83^, the authors noted antisense overlaps ranging from 50 to 2351 nt (median = 212 nt; mean = 244 nt) in all viral groups except +ssRNA viruses. The latter contains an unusual exception constituting an antisense ORF of ~1000 AA in the family *Narnaviridae*^84^. This ORF was hypothesized to be protein-coding^85^, but experimental evidence is lacking.

Almost all proposed antisense OLGs lack a native proof of the encoded protein product, arguably calling their coding potential into question. Proteomic detection of the OLG *cosA* via MS, for instance, failed, presumably due to its low expression^86^. In addition to low protein abundance, the generally small size of OLGs also hampers a proteomic proof due to an insufficient amount, or complete absence, of mass spectrometry-detectable peptides^22^. Nevertheless, protein evidence of antisense OLGs was provided in some proteomic studies^87^, but mainly attributed to a high false positive rate. Proteomic OLG evidence was found for other bacterial genera, including *Helicobacter*^65^, *Salmonella*^88^ or *Pseudomonas*^48,86^. In *P. putida,* 44 small antisense-encoded proteins were found using MS^48^. For a different species, *P. fluorescens*, nine protein-coding antisense OLGs were found using MS^86^. In the latter, eight of nine detected proteins were shorter than 200 AA; but one had a reported length of 530 AA. To our knowledge, the longest antisense OLG with proteomic evidence is a 1644 nt ORF (encoding for 548 AA), located in frame −1 in *Deinococcus radiodurans*^88^. However, up until now all prokaryotic OLGs identified via MS have lacked verification. Thus, *olg1* and *olg2* not only represent antisense OLGs of exceptional sizes across bacteria, archaea and viruses, but constitute the longest known OLGs with reliable proteomic evidence.

*olg1* and *olg2* are phylogenetically young genes under selection. For both *olg1* and *olg2*, the OLG sequence is evolving considerably faster at the AA level than the mother gene protein sequence (approximately two and 12 times faster respectively; Supplementary Table 8). This appears to have resulted in a long ORF ‘opening up’ in the recent history of *Pseudomonas* genomes for *olg1* and perhaps somewhat earlier for *olg2.* At some point, they became subject to purifying selection, as shown by depletion of stop codons, of non-synonymous changes, and of synonymous variants in the mother gene. The yet unaccounted-for evidence of translation upstream of *olg1* shown here raises the possibility of multiple start sites, which have been recently observed for many bacterial proteins^89^, including potentially for the OLG *pop*^15^. However, in our sliding window analyses of *tle3* (Fig. 5bc), we found no evidence for selection on upstream sequences.

Bioinformatic analysis of OLGs is still in its infancy. For instance, for evolutionary simulation, it would be ideal to start with the actual ancestral sequence, but ancestral-sequence reconstruction for OLGs is yet unsolved. Thus, for the simulation method, rather than introducing new biases with imperfect reconstruction, we instead followed the approach of Cassan, et al.^74^ of using a known leaf sequence in place of the root sequence. Further, choosing an outgroup with intact ORF to root the tree implicitly assumes that the ancestor of the outgroup and OLG clade contained an intact OLG, and the results are sensitive to the choice of sequence on which the tree is rooted (Supplementary Figure 8b). Here also, ancestral-sequence construction would assist with realistic simulations. Further, another limitation with existing methods is that they all use only a subset of the sequence information, e.g. ‘Frameshift’ only considers stop codons in one genome, ‘FRESCo’ only considers synonymous sites in the mother gene, and ‘OLGenie’ is restricted to the nonsynonymous mother gene sites. Future developments combining features should increase accuracy. Additional considerations such as masking out RNA secondary structures, using machine-learning methods to find subtle signatures of selection, or including sequences from metagenomes studies of different niches should improve our understanding of the evolution of OLGs and other taxonomically restricted genes.

Until recently it was thought that almost all modern genes arose through duplication and divergence from ancient genes^38^. Many taxonomically restricted genes, found only in one strain or relatively few closely related genomes, have recently been discovered. The origin of few of these “orphan” genes, however, has been explicated in molecular detail. Young OLGs have some important advantages in the study of gene evolution. In particular, the genetic context is fixed due to the presence of the mother gene. This dramatically reduces the major problems associated with false homologs and failure to detect true homologs^90,91^. The evolutionary processes involved in the initial expression and neo-functionalization of these ORFs deserve further attention. For instance, a shift in function in PA1383 appears to have involved substantial sequence change, including gain of a signal peptide. We hypothesize that during this process of positive selection on the mother gene many possible sequences were explored in the antisense −1 frame, facilitating the origin of the ORF encoding *olg2*.

Our results demonstrate that bacterial genomics after decades of advance still has additional fundamental secrets to reveal^92,93^, potentially including many more long OLGs, which were until now hiding in the shadows of known, annotated genes. These elements have not been rigorously searched for before at a whole genome level, as appropriate detection methods are still in development; and if found they are often disregarded. In this discovery of long OLGs, new research opportunities are opened for genomics, proteomics, and translatomics, as well as in the study of evolutionary novelty and bacterial gene function. These findings together shine a spotlight on the remarkable coding flexibility enabled by the redundancy in the standard genetic code.

## Methods

### Oligonucleotides and peptides

Oligonucleotides and synthetic peptides are listed in Supplementary Table 9. For Olg1, Olg2, Tle3, and PA1383 in total eighteen optimal peptides were selected for isotopically-labeled reference peptides (SpikeTidesL) purchased from JPT Peptide Technologies. Either the C-terminal lysine (Lys8) or arginine residue (Arg10) were 13C- and 15N-labeled. Isotope-labelled peptides were not purified and, thus, concentrations represent only estimates.

### Cultivation and harvest

Lysogeny broth (10 g/L tryptone, 5 g/L yeast extract, 5 g/L NaCl) was inoculated 1:100 using an overnight culture of *P. aeruginosa* PAO1 (DSM19880) and aerobically incubated (37°C, 150 rpm). After 1h, 2h, 4h, 6h, 8h, and 24h and at OD_600nm_ = 1, samples were taken by centrifugation (10 min, 12,000×g, 4°C). For transcriptomes and translatomes, cellular processes were stalled at OD_600nm_ = 1 by adding dry ice reaching 4°C. Next, cells were centrifuged (8,000×g, 4°C, 5 min) and resuspended in polysome-lysis-buffer^94^ (325 μL per 100 mL initial culture). Cells were lysed in a cell crusher with liquid nitrogen. After centrifugation as before, the supernatant was used for transcriptomes and translatomes.

### RNA isolation

Total RNA was extracted from cells (qPCR) or cell lysate (transcriptomes and translatomes) using Trizol (Thermo Fisher Scientific). For qPCR, cell pellets were resuspended in 1 mL cooled Trizol and subjected to bead beating (0.1 mm zirconia beads) using a FastPrep (3 cycles, 6.5 ms^−1^, 45 s; with 5 min incubation on ice after each cycle). Cell lysates (see above) were incubated each 5 min with first cooled Trizol and next 200 μL chloroform. After centrifugation (15 min, 12,000×g, 4°C), RNA was precipitated (500 μL 2-propanol, 1 μL glycogen, 30 min). RNA was pelleted (10 min, 12,000×g, 4°C) and washed twice with cold 70% ethanol. Air-dried RNA was dissolved in RNase-free water. Integrity was verified by agarose gel electrophoresis (1.5%, 100 V, 45 min; Carl Roth) and Bioanalyzer measurements (RNA 6000 Nano Assay, Agilent Technologies).

RNA samples were incubated with TURBO DNase (Thermo Fisher Scientific; 1 h, 60°C) and 25 U SUPERase·In RNase Inhibitor (Thermo) for removing residual DNA. After inactivation (15 mM EDTA, 10 min, 65°C), the RNA was precipitated overnight (−20°C) using ethanol, 3 M sodium acetate, and glycogen (690, 27.6, and 1 μL, respectively). Precipitated RNA was pelleted, washed, dried and dissolved as before. DNA absence was confirmed with PCR using *Taq* polymerase (NEB) with primer 1 & 2.

### cDNA synthesis for PCR

RNA (500 μg) was reverse-transcribed using SuperScript III Reverse Transcriptase (Thermo Fisher Scientific). Random nonamer (50 pmol; Sigma Aldrich) or 10 pmol primer 3 were used for reverse transcription of *gyrA* (reference gene) or *olg1*, respectively, in the presence of 20 U SUPERase·In RNase Inhibitor. Samples without reverse transcriptase served as negative controls. For transcriptional termination sites of *olg1,* reverse transcription was performed with primer 4 or 5. Of the latter, 1 μL cDNA was used in a 30-cycle PCR using Taq DNA Polymerase (NEB) with primer 4 & 6 or 5 & 6. For *olg2*, RNA was reversed transcribed using primer 7. Subsequently, 1 μL cDNA was used in a 30-cycle PCR using Q5 DNA Polymerase (NEB) with primer 8 & 9 or 10 & 11. cDNA of *olg1* was additionally used testing for alternative start sites by PCR using the primer 8 & 15, 8 & 16, 8 & 17, and 8 &18. Primer functionalities were verified with genomic DNA before (not shown).

### Quantitative PCR

Expression levels of *olg1, olg2* and *gyrA* were quantified by qPCR. Each 20-μL reaction contained 10 μL SsoAdvanced Universal SYBR Green Supermix (Bio-Rad Laboratories), 500 nM forward and reverse primer, and 1 μL cDNA (or water for “No Template Control”). For *gyrA* and *olg1*, primer 12 & 13 and 6 & 14 were used, respectively. Cycling was as follows: 95°C for 30 s; 40 cycles of 95°C for 15 s and 60°C for 30 s. A melt curve analysis (65 to 95°C with 0.5°C increments) confirmed the correct product. Each reaction was conducted in three biological and technical replicates. Data were analysed using the ΔΔCt method^95^. Significance was evaluated with a two-tailed Welch two-sample t-test (*p*-value ≤0.05).

### Transcriptome sequencing

rRNA was depleted from total RNA (of 200 μL cell-extract, DNase treated, as above) using the *P. aeruginosa*-specific riboPOOL kit (version v1-5, siTOOLs Biotech) followed by RNA precipitation and DNase digestion of the probes. One μg depleted RNA was fragmented (Ultrasonicator system S220, Covaris; 175 W, 10% duty cycle, 200 cycles for 180 s), dephosphorylated (Antarctic phosphatase, NEB), and phosphorylated (T4 Polynucleotide Kinase, NEB). Fragments were purified after each step using the miRNeasy Mini Kit (Qiagen). Finally, volume was reduced to 5 μL in a Speedvac concentrator (Eppendorf) and sequencing libraries were prepared using the TruSeq Small RNA Library Prep Kit (Illumina). cDNA concentration and length were measured using a Qubit (dsDNA HS Assay Kit, Thermo Fisher) and Bioanalyzer (High Sensitivity DNA Kit, Agilent). Libraries were diluted to 2 nM in 10 μL 10 mM Tris-HCl (pH 8.5) and sequenced on a HiSeq2500 (Illumina) using a v2 Rapid SR50 cartridge (Illumina) for two biological replicates.

### Ribosome Profiling

Translatome sequencing was conducted as described^96^ with a few modifications. Briefly, 25 absorption units of the cell lysate were incubated (1 h, 25°C, shaking at 850 rpm) with 62.5 U MNase (Thermo Fisher), 18.75 U RNase R (Lucigen), 4.375 U RNase T (NEB), 1.875 U XRN-1 (NEB), 1 mM CaCl_2_, and 1×Buffer 4 (NEB). The reaction was stopped (6 mM EGTA, 50 U SUPERase·In, 10 min). Monosomes were isolated by sucrose density gradient centrifugation (104,000×g, 4°C, 3 h) followed by RNA isolation and DNase treatment as described. Ribosomal footprints were size selected using a 16% denaturing urea polyacrylamide gel (Carl Roth; 200 V, 1.5 h). After staining (SYBR Gold, Invitrogen), ribosomal footprints (19 – 27 nt) were excised. Gel pieces were crushed in gel breaker tubes (15,700 ×g, 2 min). Gel debris was incubated overnight in extraction buffer (300 mM NaOAc pH 5.5, 1 mM EDTA, 0.1 U/μL SUPERase·In). After centrifugation (2 min, 9,300×g, RT) in 0.22-μm pore cellulose-acetate filter tubes, footprints were precipitated with ethanol and transformed in a sequencing library as above for two biological replicates.

### Cell lysis and protein digest for mass spectrometry

Cells were lysed in 100 μL absolute TFA (Sigma-Aldrich; 5 min, 55°C, shaking at 1,000 rpm) and neutralized with 900 μL 2 M Tris^97^. Protein concentration was determined using Bradford reagent (B6916, Sigma-Aldrich). For offline high pH reversed-phase (hpH RP) fractionation and for targeted proteomics, 75 μg and 20 μg of total protein amount were reduced and alkylated (10 mM TCEP, 55 mM CAA; 5 min, 95°C), respectively. Water-diluted samples (1:1) were subjected to proteolysis with trypsin (enzyme to protein ratio 1:50, 30°C, overnight, shaking at 400 rpm) and then stopped (3% formic acid, FA).

### Offline high pH reversed-phase fractionation for full proteome analysis

Three discs of Empore C18 (3M) material were packed in 200-μL pipette tips. The resulting desalting columns were conditioned (100% acetonitrile, ACN) and equilibrated (40% ACN/0.1% FA) followed by 2% ACN/0.1% FA. Peptides of the 75-μg protein digest were loaded, washed (2% ACN/0.1% FA) and eluted (40% ACN/0.1% FA). Next, peptides were fractionated using an Agilent 1100 series HPLC system operating a XBridge BEH130 C18 3.5 μm 2.1 × 250 mm column (Waters) at a flow rate of 200 μL/min. Buffer A was 25 mM ammonium bicarbonate (pH 8.0), buffer B was 80% ACN. Fractions were collected every minute into a 96 well plate. Peptides were separated by a linear gradient from 4% to 32% buffer B over 45 min, followed by a gradient from 32% to 85% buffer B over 6 min. Samples were collected in 30 s steps between minute 3 and 51. The solvent was evaporated and samples were redissolved in 2% ACN/0.1% FA.

### High pH reversed-phase fractionation for targeted proteomics

C18-packed 200-μl tips (see above) were loaded with peptides from the 20 μg digest. A pH switch was performed using 25 mM ammonium formate (pH 10) and varying ACN concentrations for each of six fractions. ACN was added at concentrations of 0, 5, 10, 15, 25, and 50%, respectively. Fraction 1 and 5 and fraction 2 and 6 were combined. The solvent was each evaporated (1+5, 2+6, 3, and 4), and samples were dissolved in 2% ACN/0.1% FA.

### LC-MS/MS measurements - Full proteomes

Peptides were analysed on a Dionex Ultimate 3000 RSLCnano system coupled to a Fusion Lumos Tribrid mass spectrometer (Thermo Fisher Scientific). For full proteome analyses, 0.5 μg of peptides were delivered to a trap column (ReproSil-pur C18-AQ, 5 μm, 20 mm × 75 μm, Dr. Maisch, self-packed) at a flow rate of 5 μL/min of solvent A (HPLC-grade water with 0.1% FA). After loading (10 min), peptides were transferred to an analytical column (ReproSil Gold C18-AQ, 3 μm, 450 mm × 75 μm, Dr. Maisch, self-packed) and separated using a 50-min linear gradient from 4% to 32% of solvent B (ACN/0.1% FA/5% dimethyl sulfoxide, DMSO) in solvent A (HPLC-grade water with 0.1% FA/5% DMSO) at 300 nL/min flow rate. Both solvents contain DMSO boosting MS intensity. The Fusion Lumos Tribrid mass spectrometer was operated in data-dependent acquisition (DDA) and positive ionization mode. MS1 spectra (360–1300 m/z) were recorded at a resolution of 60,000 using an automatic gain control (AGC) target value of 4×10^5^ and maximum injection time (MaxIT) of 50 ms. Up to 20 peptide precursors were selected for fragmentation in case of the full proteome analyses. Only precursors with charge state 2 to 6 were selected and dynamic exclusion of 20 s was enabled. Peptide fragmentation was performed using higher energy collision induced dissociation (HCD) and a normalized collision energy (NCE) of 30%. The precursor isolation window width was set to 1.3 m/z. MS2 spectra were acquired in the orbitrap with a resolution of 15.000 and an AGC target value of 5×10^4^. MaxIT was set to 22 ms.

### Mass spectrometric data analysis - Full proteomes

Peptide identification and quantification was performed using MaxQuant^98^ (v1.6.3.4) with Andromeda^99^. MS2 spectra were searched against the RefSeq^100^ file downloaded for *P. aeruginosa* PAO1 (GCF_000006765.1_ASM676v1_protein.faa, 5,572 reviewed entries, 7 Feburary 2020), supplemented with common contaminants (by MaxQuant) and Olg1 and Olg2 AA sequences. Trypsin/P was specified as proteolytic enzyme. Precursor tolerance was set to 4.5 ppm and fragment ion tolerance to 20 ppm. Results were adjusted to 1% FDR on peptide spectrum match level and protein level employing a target-decoy approach using reversed protein sequences. Minimal peptide length was defined as 7 AA; the “match-between-run” function disabled. For full proteome analyses, carbamidomethylated cysteine was set as fixed and oxidation of methionine and N-terminal protein acetylation as variable modifications. Correlation scores (dot product) between experimental and predicted spectra were calculated via Skyline daily (64-bit, v20.1.9.234)^101^ that supports Prosit^68^ spectra predictions. For data analysis, protein intensities and iBAQ^102^ values were calculated.

### Targeted LC-MS/MS measurements

Targeted measurements using Parallel Reaction Monitoring (PRM) were performed with a 50-min linear gradient on a Dionex Ultimate 3000 RSLCnano system coupled to a Q-Exactive HF-X mass spectrometer (Thermo Fisher Scientific). The spectrometer was operated in PRM and positive ionization mode. MS1 spectra (360–1300 m/z) were recorded at a resolution of 60,000 using an AGC target value of 3×10^6^ and a MaxIT of 100 ms. Targeted MS2 spectra were acquired at 60,000 resolution with a fixed first mass of 100 m/z, after HCD with 26% NCE, and using an AGC target value of 1×10^6^, a MaxIT of 118 ms and an isolation window of 0.7 m/z. For the PRM analysis of the growth phase samples, 18 OLG and mother gene peptides plus 12 retention time reference peptides (subset of Procal peptides synthesized by JPT^103^) were targeted within a single PRM run and with a 5 min scheduled retention time window. The cycle time was ~2.1 s, which leads to ~10 data points per chromatographic peak.

### Selection and validation of target peptides

Isotope-labelled internal reference peptides were used for confident identification and quantification. Peptide selections were based on results of DDA measurements of the deep proteome at OD_600nm_ = 1. Peptides were selected based on intensity, location within the protein, Andromeda score, excluding modification, and charge state. All isotopically-labeled synthetic peptides were pooled and targeted proteomic measurements (PRM) showed confident detection of all 18 peptides (MaxQuant score >90, Supplementary Table 10). Skyline-daily^101^ was used to build an experimental spectral library from the generated PRM data.

### Targeted mass spectrometric data analysis

PRM data was analysed using Skyline-daily^101^. Peak integration, transition interferences and integration boundaries were reviewed manually, considering four to six transitions per peptide. To discriminate between positive and negative peptide detection, filtering according to correlation of fragment ion intensities between the endogenous (light) and the spike-in (heavy) peptides was applied (“Library Dot Product” ≥0.8). Additionally, a correlation of fragment ion intensities between the light and heavy peptide (“DotProductLightToHeavy” of >0.9) and a mass accuracy of below ±20 ppm (“Average Mass Error PPM”) was required. Total protein intensity was computed by summing up all light peptide intensities detected positive in each sample (Supplementary Fig. 6b). Uniqueness of the peptides was assessed against the RefSeq database for *P. aeruginosa* PAO1.

### Bioinformatic analyses

Putative σ70 promoters within a 300-nt region upstream of the start codon were predicted by BPROM^58^ with minimum LDF scores of 0.2.

Shine-Dalgarno sequence identification was performed as described^61^ within a region of 30 nt upstream of the start codon and a minimum free energy (*ΔG*_*SD*_) threshold of −2.9 kcal/mol.

To predict ρ-independent terminators, a 300-nt region downstream of the respective stop codon was analysed using FindTerm^58^ with an threshold of −3. Predicted terminator regions were read in non-overlapping sliding windows of 30 nt and folded with Mfold^104^, identifying stem loops.

FASTQ files were processed using a custom perl script. FastQC^105^ was used to assess raw read quality and adapter sequences were trimmed with fastp^106^. Trimmed reads were aligned to the reference (GCF_000006765.1_ASM676v1) with Bowtie2 v2.2.6^107^ using “--very-sensitive end-to-end” with a seed length of 17 nt. Reads mapping to rRNAs and tRNAs were filtered with SAMTools^108^ and BEDTools^109^. Remaining reads were normalized to gene length and sequencing depth (RPKM: “reads per kilobase per million mapped reads”). For each genomic nucleotide position, reads per million mapped reads (RPM) were calculated, averaged over biological replicates and visualized with pyGenomeTracks^110^. Ribosome coverage values^111^ (RCV) were calculated by dividing the RPKM of the translatome by the RPKM of the transcriptome for evaluating ‘translatability’.

Significant changes in translation were determined between the published RiboSeq datasets “M9+n-alkane” and “M9+glycerol”^69^. Read counts were scaled to the smallest library size and differential expression analysis was performed using an exact test implemented in edgeR^112^.

Gene prediction was performed with Prodigal^62^ using default parameters. In order to detect overlapping ORFs, all possible start codons within and in the upstream-vicinity of the coding regions for *tle3* and PA1383 were masked by N (any nucleotide). Further, protein-coding ORFs were predicted based on RiboSeq data using DeepRibo^66^ (default settings).

The Basic Local Alignment Search Tool (blast)^113^ was used to find homologous nucleotide (blastn) and protein sequences (blastp).

### Phylostratigraphy – taxonomic distribution

Homologs of *tle3* and PA1383 (NC_002516.2) were detected using BLASTp in annotated proteins from genomes in Pseudomonadales, and from the Identical Protein Groups database using the Entrez Programming Utilities^114^. Sequences from MAG collections were added using Diamond blastp^115^ finding homologs within genomes annotated as being within Pseudomonadales. The combined collection was downsampled to unique nucleotide sequences with a length of ≥50% compared to *tle3* or PA1383 in the reference strain PAO1.

Protein sequences were aligned using QuickProbs2^116^, and nucleotide sequences subsequently converted to codon alignments using Pal2Nal^117^. Maximum likelihood trees of *tle3* and PA1383 alignments were calculated using IQ-TREE^118,119^ with default settings and 1000 bootstrap iterations. Trees were downsampled to 20 genomes using Treemmer^120^, and edited using Newick Utilities^121^ for OLG-region visualisation using a Python script written with ETE3 packages^122^, and subsequently also used for pairwise ‘OLGenie’ comparisons^30^.

### ORF length and stop codon analyses

Excess overlapping ORF lengths were tested for using ‘Frameshift’ ^73^, modified to run from the Unix command line, as well as to print the scores obtained for each method (codon permutation and synonymous codon mutation), and to print the simulated ORF lengths.

An evolutionary simulation method^74^ was re-implemented using IQ-TREE and Pyvolve^75^, with sequence evolution simulated along trees calculated from the OLG-containing genomes and a selected outgroup sequence without stop codons in the OLG loci. In the original study, an outgroup with an intact ORF was used for rooting the tree^74^. A sequence from *P. prosekii*, the only intact homolog outside the OLG clade of *P. aeruginosa*, was chosen as outgroup for *olg1*. For *olg2*, multiple non-*P. aeruginosa* intact ORFs were available. A more distant outgroup was chosen as tests of purifying selection described below suggest more taxonomically widespread functionality. Omega values (approximately equivalent to dN/dS) of 0.5 for both genes were chosen based on alignments of the two mother genes, using results from ‘OLGenie’. The empirical codon model (ECM) was used with default parameters except for the omega values specified.

### Codon-position constraint analyses

Constraints in synonymous sites of *tle3* and PA1383 were assessed using ‘FRESCo’^28^. Approximate maximum-likelihood nucleotide trees were calculated using FastTree^123^ for the full sets of “OLG” and “non-OLG” genomes, and ‘FRESCo’ was run on codon alignments (described above) with a sliding window size (50 codons).

Constraint on non-synonymous codon changes in the OLGs was assessed using ‘OLGenie’^30^. Analysis for each mother-gene codon alignment (created using PAL2NAL, described above) of OLG and non-OLG genomes was conducted with standard settings. Sliding window analyses of 50 codons were conducted using a minimum number of defined codons of 2. Pairwise whole-gene comparisons of *olg1* and *olg2* were conducted using standard settings, and a custom Bash script producing a pairwise matrix.

## Data availability

Further data are available in the Supplementary Information. Sequencing data have been deposited in Sequence Read Archive (NCBI) with the accession [pending]. The proteomics raw data, MaxQuant search results and used protein sequence databases have been deposited with the ProteomeXchange Consortium via the PRIDE partner repository^124^ and can be accessed using the data set identifier PXD023992. All targeted proteomic raw data and Skyline analysis files have been deposited to Panorama Public^125^ [pending]. Scripts for evolutionary and taxonomic analyses are available at https://github.com/ZacharyArdern/Pseudomonas_long_OLGs.

## Acknowledgements

We thank Romy Wecko, Verena Breitner, Lara Wanner, Hermine Kienberger, and Franziska Hackbarth for technical assistance, and Christopher Huptas for bioinformatic support. We also thank Siddhanth Rao for assistance with scripts for the use of ‘FRESCo’, and Chase Nelson and April Wei for helpful comments on the manuscript.

## Author contributions

MK conducted cultivation and sequencing experiments and drafted the manuscript. ZA designed and performed evolutionary analyses. MA conducted mass spectrometry experiments with the help of CL. SS and KN conceived the study. SS provided funding. All authors contributed in writing.

## Competing interests

The authors declare no competing interests.

## Additional information

Correspondence and requests concerning evolutionary analyses should be addressed to Z.A.; concerning biological experiments to K.N.

## Notes

https://github.com/ZacharyArdern/Pseudomonas_long_OLGs

### Competing Interest Statement

The authors have declared no competing interest.

## References

1 Barrell, B. G., Air, G. M. & Hutchison, C. A., 3rd. Overlapping genes in bacteriophage phiX174. Nature 264, 34–41, doi:10.1038/264034a0 (1976).

2 Delcher, A. L., Bratke, K. A., Powers, E. C. & Salzberg, S. L. Identifying bacterial genes and endosymbiont DNA with Glimmer. Bioinformatics (Oxford, England) 23, 673–679, doi:10.1093/bioinformatics/btm009 (2007).

3 Warren, A. S., Archuleta, J., Feng, W.-c. & Setubal, J. C. Missing genes in the annotation of prokaryotic genomes. BMC Bioinformatics 11, 131, doi:10.1186/1471-2105-11-131 (2010).

4 Yooseph, S. et al. The Sorcerer II Global Ocean Sampling expedition: expanding the universe of protein families. PLoS Biology 5, e16 (2007).

5 Salzberg, S. L., Delcher, A. L., Kasif, S. & White, O. Microbial gene identification using interpolated Markov models. Nucleic Acids Res 26, 544–548, doi:10.1093/nar/26.2.544 (1998).

6 Jensen, K. T. et al. Novel overlapping coding sequences in *Chlamydia trachomatis*. FEMS Microbiology Letters 265, 106–117, doi:10.1111/j.1574-6968.2006.00480.x (2006).

7 Baek, J., Lee, J., Yoon, K. & Lee, H. Identification of Unannotated Small Genes in *Salmonella*. G3 (Bethesda) 7, 983–989, doi:10.1534/g3.116.036939 (2017).

8 Weaver, J., Mohammad, F., Buskirk, A. R. & Storz, G. Identifying Small Proteins by Ribosome Profiling with Stalled Initiation Complexes. mBio 10, e02819–02818, doi:10.1128/mBio.02819-18 (2019).

9 Behrens, M., Sheikh, J. & Nataro, J. P. Regulation of the overlapping pic/set locus in *Shigella flexneri* and enteroaggregative *Escherichia coli*. Infect Immun 70, 2915–2925, doi:10.1128/iai.70.6.2915-2925.2002 (2002).

10 Fellner, L. et al. Phenotype of htgA (mbiA), a recently evolved orphan gene of *Escherichia coli* and *Shigella*, completely overlapping in antisense to yaaW. FEMS Microbiology Letters 350, 57–64, doi:10.1111/1574-6968.12288 (2014).

11 Fellner, L. et al. Evidence for the recent origin of a bacterial protein-coding, overlapping orphan gene by evolutionary overprinting. BMC Evol Biol 15, 283–283, doi:10.1186/s12862-015-0558-z (2015).

12 Hücker, S. M., Vanderhaeghen, S., Abellan-Schneyder, I., Scherer, S. & Neuhaus, K. The Novel Anaerobiosis-Responsive Overlapping Gene ano Is Overlapping Antisense to the Annotated Gene ECs2385 of *Escherichia coli* O157:H7 Sakai. Frontiers in microbiology 9, 931–931, doi:10.3389/fmicb.2018.00931 (2018).

13 Hücker, S. M. et al. A novel short L-arginine responsive protein-coding gene (laoB) antiparallel overlapping to a CadC-like transcriptional regulator in *Escherichia coli* O157:H7 Sakai originated by overprinting. BMC Evol Biol 18, 21–21, doi:10.1186/s12862-018-1134-0 (2018).

14 Vanderhaeghen, S., Zehentner, B., Scherer, S., Neuhaus, K. & Ardern, Z. The novel EHEC gene asa overlaps the TEGT transporter gene in antisense and is regulated by NaCl and growth phase. Scientific reports 8, 17875 (2018).

15 Zehentner, B., Ardern, Z., Kreitmeier, M., Scherer, S. & Neuhaus, K. A Novel pH-Regulated, Unusual 603 bp Overlapping Protein Coding Gene pop Is Encoded Antisense to ompA in *Escherichia coli* O157:H7 (EHEC). Frontiers in Microbiology 11, doi:10.3389/fmicb.2020.00377 (2020).

16 Zehentner, B., Ardern, Z., Kreitmeier, M., Scherer, S. & Neuhaus, K. Evidence for Numerous Embedded Antisense Overlapping Genes in Diverse *E. coli* Strains. bioRxiv, 2020.2011.2018.388249, doi:10.1101/2020.11.18.388249 (2020).

17 Smith, C. et al. Pervasive Translation in *Mycobacterium tuberculosis*. bioRxiv, 665208, doi:10.1101/665208 (2019).

18 Filiatrault, M. J. et al. Transcriptome Analysis of *Pseudomonas syringae* Identifies New Genes, Noncoding RNAs, and Antisense Activity. Journal of Bacteriology 192, 2359, doi:10.1128/JB.01445-09 (2010).

19 Gelsinger, D. R. et al. Ribosome profiling in archaea reveals leaderless translation, novel translational initiation sites, and ribosome pausing at single codon resolution. Nucleic Acids Res 48, 5201–5216, doi:10.1093/nar/gkaa304 (2020).

20 Loughran, G. et al. Unusually efficient CUG initiation of an overlapping reading frame in POLG mRNA yields novel protein POLGARF. Proceedings of the National Academy of Sciences 117, 24936, doi:10.1073/pnas.2001433117 (2020).

21 Khan, Y. A. et al. Evidence for a novel overlapping coding sequence in POLG initiated at a CUG start codon. BMC Genetics 21, 25, doi:10.1186/s12863-020-0828-7 (2020).

22 Petruschke, H., Anders, J., Stadler, P. F., Jehmlich, N. & von Bergen, M. Enrichment and identification of small proteins in a simplified human gut microbiome. Journal of Proteomics 213, 103604 (2020).

23 Ardern, Z., Neuhaus, K. & Scherer, S. Are Antisense Proteins in Prokaryotes Functional? Frontiers in molecular biosciences 7, 187–187, doi:10.3389/fmolb.2020.00187 (2020).

24 Ingolia, N. T., Ghaemmaghami, S., Newman, J. R. S. & Weissman, J. S. Genome-wide analysis in vivo of translation with nucleotide resolution using ribosome profiling. Science 324, 218–223, doi:10.1126/science.1168978 (2009).

25 Fremin, B. J. & Bhatt, A. S. Structured RNA Contaminants in Bacterial Ribo-Seq. Msphere 5 (2020).

26 Sabath, N., Landan, G. & Graur, D. A method for the simultaneous estimation of selection intensities in overlapping genes. PLoS One 3, e3996 (2008).

27 Firth, A. E. Mapping overlapping functional elements embedded within the protein-coding regions of RNA viruses. Nucleic acids research 42, 12425–12439 (2014).

28 Sealfon, R. S. et al. FRESCo: finding regions of excess synonymous constraint in diverse viruses. Genome biology 16, 38 (2015).

29 Wei, X. & Zhang, J. A simple method for estimating the strength of natural selection on overlapping genes. Genome biology and evolution 7, 381–390 (2015).

30 Nelson, C. W., Ardern, Z. & Wei, X. OLGenie: Estimating Natural Selection to Predict Functional Overlapping Genes. Molecular Biology and Evolution, doi:10.1093/molbev/msaa087 (2020).

31 Meydan, S. et al. Retapamulin-assisted ribosome profiling reveals the alternative bacterial proteome. Molecular cell 74, 481–493. e486 (2019).

32 Mir, K., Neuhaus, K., Scherer, S., Bossert, M. & Schober, S. Predicting statistical properties of open reading frames in bacterial genomes. PLoS One 7 (2012).

33 Koskella, B. & Brockhurst, M. A. Bacteria–phage coevolution as a driver of ecological and evolutionary processes in microbial communities. FEMS microbiology reviews 38, 916–931 (2014).

34 Sander, C. & Schulz, G. E. Degeneracy of the information contained in amino acid sequences: evidence from overlaid genes. Journal of molecular evolution 13, 245–252 (1979).

35 Miyata, T. & Yasunaga, T. Evolution of overlapping genes. Nature 272, 532 (1978).

36 Yockey, H. P. Do overlapping genes violate molecular biology and the theory of evolution? J Theor Biol 80, 21–26 (1979).

37 Portelli, G. The relations between the precodons of overlapping genes. Journal of theoretical biology 95, 345–350 (1982).

38 Ohno, S. Evolution by gene duplication. (Allen & Unwin; Springer-Verlag, 1970).

39 Grassé, P. P. in Evolution of living organisms: evidence for a new theory of transformation 231–237 (Academic Press, 1977).

40 Keese, P. K. & Gibbs, A. Origins of genes: "big bang" or continuous creation? Proc Natl Acad Sci U S A 89, 9489–9493, doi:10.1073/pnas.89.20.9489 (1992).

41 Bartonek, L., Braun, D. & Zagrovic, B. Frameshifting preserves key physicochemical properties of proteins. Proceedings of the National Academy of Sciences 117, 5907, doi:10.1073/pnas.1911203117 (2020).

42 Nelson, C. W. et al. Dynamically evolving novel overlapping gene as a factor in the SARS-CoV-2 pandemic. Elife 9, e59633, doi:10.7554/eLife.59633 (2020).

43 Konecny, J., Eckert, M., Schöniger, M. & Hofacker, G. L. Neutral adaptation of the genetic code to double-strand coding. Journal of Molecular Evolution 36, 407, doi:10.1007/BF02406718 (1993).

44 Tautz, D. & Domazet-Lošo, T. The evolutionary origin of orphan genes. Nature Reviews Genetics 12, 692–702 (2011).

45 Storz, G., Wolf, Y. I. & Ramamurthi, K. S. Small proteins can no longer be ignored. Annual review of biochemistry 83, 753–777, doi:10.1146/annurev-biochem-070611-102400 (2014).

46 Silby, M. W. & Levy, S. B. Overlapping protein-encoding genes in *Pseudomonas fluorescens* Pf0-1. PLoS Genet 4, e1000094–e1000094, doi:10.1371/journal.pgen.1000094 (2008).

47 Silby, M. W. & Levy, S. B. Use of in vivo expression technology to identify genes important in growth and survival of *Pseudomonas fluorescens* Pf0-1 in soil: discovery of expressed sequences with novel genetic organization. J Bacteriol 186, 7411–7419, doi:10.1128/jb.186.21.7411-7419.2004 (2004).

48 Yang, X., Jensen, S. I., Wulff, T., Harrison, S. J. & Long, K. S. Identification and validation of novel small proteins in *Pseudomonas putida*. Environmental Microbiology Reports 8, 966–974, doi:10.1111/1758-2229.12473 (2016).

49 Crone, S. et al. The environmental occurrence of *Pseudomonas aeruginosa*. APMIS 128, 220–231, doi:https://doi.org/10.1111/apm.13010 (2020).

50 Kerr, K. G. & Snelling, A. M. *Pseudomonas aeruginosa*: a formidable and ever-present adversary. Journal of Hospital Infection 73, 338–344, doi:10.1016/j.jhin.2009.04.020 (2009).

51 Weinstein, R. A., Gaynes, R., Edwards, J. R. & National Nosocomial Infections Surveillance, S. Overview of Nosocomial Infections Caused by Gram-Negative Bacilli. Clinical Infectious Diseases 41, 848–854, doi:10.1086/432803 (2005).

52 Livermore, D. M. Has the era of untreatable infections arrived? J Antimicrob Chemother 64 Suppl 1, i29–36, doi:10.1093/jac/dkp255 (2009).

53 Bassetti, M., Vena, A., Croxatto, A., Righi, E. & Guery, B. How to manage *Pseudomonas aeruginosa* infections. Drugs Context 7, 212527–212527, doi:10.7573/dic.212527 (2018).

54 Berni, B., Soscia, C., Djermoun, S., Ize, B. & Bleves, S. A Type VI Secretion System Trans-Kingdom Effector Is Required for the Delivery of a Novel Antibacterial Toxin in *Pseudomonas aeruginosa*. Front Microbiol 10, 1218, doi:10.3389/fmicb.2019.01218 (2019).

55 Lewenza, S., Gardy, J. L., Brinkman, F. S. L. & Hancock, R. E. W. Genome-wide identification of *Pseudomonas aeruginosa* exported proteins using a consensus computational strategy combined with a laboratory-based PhoA fusion screen. Genome research 15, 321–329, doi:10.1101/gr.3257305 (2005).

56 Crespo, A., Pedraz, L. & Torrents, E. Function of the *Pseudomonas aeruginosa* NrdR Transcription Factor: Global Transcriptomic Analysis and Its Role on Ribonucleotide Reductase Gene Expression. PLOS ONE 10, e0123571, doi:10.1371/journal.pone.0123571 (2015).

57 Lippa, A. M., Gebhardt, M. J. & Dove, S. L. H-NS-like proteins in *Pseudomonas aeruginosa* coordinately silence intragenic transcription. Mol Microbiol, doi:10.1111/mmi.14656 (2020).

58 Solovyev, V. & Salamov, A. Automatic annotation of microbial genomes and metagenomic sequences. Metagenomics and its applications in agriculture, biomedicine and environmental studies, 61–78 (2011).

59 Wurtzel, O. et al. The single-nucleotide resolution transcriptome of *Pseudomonas aeruginosa* grown in body temperature. PLoS pathogens 8, e1002945–e1002945, doi:10.1371/journal.ppat.1002945 (2012).

60 Filiatrault, M. J. et al. Genome-wide identification of transcriptional start sites in the plant pathogen *Pseudomonas syringae* pv. tomato str. DC3000. PloS one 6, e29335–e29335, doi:10.1371/journal.pone.0029335 (2011).

61 Ma, J., Campbell, A. & Karlin, S. Correlations between Shine-Dalgarno sequences and gene features such as predicted expression levels and operon structures. Journal of bacteriology 184, 5733–5745, doi:10.1128/jb.184.20.5733-5745.2002 (2002).

62 Hyatt, D. et al. Prodigal: prokaryotic gene recognition and translation initiation site identification. BMC Bioinformatics 11, 119–119, doi:10.1186/1471-2105-11-119 (2010).

63 Eckweiler, D. & Häussler, S. Antisense transcription in *Pseudomonas aeruginosa*. Microbiology (Reading) 164, 889–895, doi:10.1099/mic.0.000664 (2018).

64 Dornenburg, J. E., Devita, A. M., Palumbo, M. J. & Wade, J. T. Widespread antisense transcription in *Escherichia coli*. mBio 1, doi:10.1128/mBio.00024-10 (2010).

65 Friedman, R. C. et al. Common and phylogenetically widespread coding for peptides by bacterial small RNAs. BMC genomics 18, 553–553, doi:10.1186/s12864-017-3932-y (2017).

66 Clauwaert, J., Menschaert, G. & Waegeman, W. DeepRibo: a neural network for precise gene annotation of prokaryotes by combining ribosome profiling signal and binding site patterns. Nucleic Acids Res 47, e36–e36, doi:10.1093/nar/gkz061 (2019).

67 Neuhaus, K. et al. Translatomics combined with transcriptomics and proteomics reveals novel functional, recently evolved orphan genes in *Escherichia coli* O157: H7 (EHEC). BMC Genomics 17, 133 (2016).

68 Gessulat, S. et al. Prosit: proteome-wide prediction of peptide tandem mass spectra by deep learning. Nat Methods 16, 509–518, doi:10.1038/s41592-019-0426-7 (2019).

69 Grady, S. L. et al. A comprehensive multi-omics approach uncovers adaptations for growth and survival of *Pseudomonas aeruginosa* on n-alkanes. BMC Genomics 18, 334, doi:10.1186/s12864-017-3708-4 (2017).

70 Almeida, A. et al. A unified catalog of 204,938 reference genomes from the human gut microbiome. Nature Biotechnology, doi:10.1038/s41587-020-0603-3 (2020).

71 Nayfach, S., Shi, Z. J., Seshadri, R., Pollard, K. S. & Kyrpides, N. C. New insights from uncultivated genomes of the global human gut microbiome. Nature 568, 505–510, doi:10.1038/s41586-019-1058-x (2019).

72 Cooper, H. B. & Gardner, P. P. Features of Functional Human Genes. bioRxiv, 2020.2010.2010.334193, doi:10.1101/2020.10.10.334193 (2020).

73 Schlub, T. E., Buchmann, J. P. & Holmes, E. C. A Simple Method to Detect Candidate Overlapping Genes in Viruses Using Single Genome Sequences. Molecular Biology and Evolution 35, 2572–2581, doi:10.1093/molbev/msy155 (2018).

74 Cassan, E., Arigon-Chifolleau, A. M., Mesnard, J. M., Gross, A. & Gascuel, O. Concomitant emergence of the antisense protein gene of HIV-1 and of the pandemic. Proceedings of the National Academy of Sciences of the United States of America 113, 11537–11542, doi:10.1073/pnas.1605739113 (2016).

75 Spielman, S. J. & Wilke, C. O. Pyvolve: a flexible Python module for simulating sequences along phylogenies. PloS one 10, e0139047 (2015).

76 Chirico, N., Vianelli, A. & Belshaw, R. Why genes overlap in viruses. Proc Biol Sci 277, 3809–3817, doi:10.1098/rspb.2010.1052 (2010).

77 Lebre, S. & Gascuel, O. The combinatorics of overlapping genes. J Theor Biol 415, 90–101, doi:10.1016/j.jtbi.2016.09.018 (2017).

78 Pallejà, A., Harrington, E. D. & Bork, P. Large gene overlaps in prokaryotic genomes: result of functional constraints or mispredictions? BMC Genomics 9, 335, doi:10.1186/1471-2164-9-335 (2008).

79 Russell, A. B. et al. Diverse type VI secretion phospholipases are functionally plastic antibacterial effectors. Nature 496, 508–512, doi:10.1038/nature12074 (2013).

80 Lynch, M. & Marinov, G. K. The bioenergetic costs of a gene. Proceedings of the National Academy of Sciences 112, 15690, doi:10.1073/pnas.1514974112 (2015).

81 Meydan, S. et al. Retapamulin-Assisted Ribosome Profiling Reveals the Alternative Bacterial Proteome. Mol Cell 74, 481–493.e486, doi:10.1016/j.molcel.2019.02.017 (2019).

82 Tunca, S., Barreiro, C., Coque, J. J. & Martín, J. F. Two overlapping antiparallel genes encoding the iron regulator DmdR1 and the Adm proteins control siderophore and antibiotic biosynthesis in *Streptomyces coelicolor* A3(2). Febs j 276, 4814–4827, doi:10.1111/j.1742-4658.2009.07182.x (2009).

83 Schlub, T. E. & Holmes, E. C. Properties and abundance of overlapping genes in viruses. Virus Evolution 6, doi:10.1093/ve/veaa009 (2020).

84 DeRisi, J. L. et al. An exploration of ambigrammatic sequences in narnaviruses. Sci Rep 9, 17982, doi:10.1038/s41598-019-54181-3 (2019).

85 Dinan, A. M., Lukhovitskaya, N. I., Olendraite, I. & Firth, A. E. A case for a negative-strand coding sequence in a group of positive-sense RNA viruses. Virus Evol 6, veaa007, doi:10.1093/ve/veaa007 (2020).

86 Kim, W. et al. Proteomic Detection of Non-Annotated Protein-Coding Genes in *Pseudomonas fluorescens* Pf0-1. PLOS ONE 4, e8455, doi:10.1371/journal.pone.0008455 (2009).

87 Venter, E., Smith, R. D. & Payne, S. H. Proteogenomic Analysis of Bacteria and Archaea: A 46 Organism Case Study. PLOS ONE 6, e27587, doi:10.1371/journal.pone.0027587 (2011).

88 Willems, P., Fijalkowski, I. & Van Damme, P. Lost and Found: Re-searching and Re-scoring Proteomics Data Aids Genome Annotation and Improves Proteome Coverage. mSystems 5, e00833–00820, doi:10.1128/mSystems.00833-20 (2020).

89 Fijalkowska, D., Fijalkowski, I., Willems, P. & Van Damme, P. Bacterial riboproteogenomics: the era of N-terminal proteoform existence revealed. FEMS Microbiology Reviews (2020).

90 Weisman, C. M., Murray, A. W. & Eddy, S. R. Many, but not all, lineage-specific genes can be explained by homology detection failure. PLOS Biology 18, e3000862, doi:10.1371/journal.pbio.3000862 (2020).

91 Vakirlis, N., Carvunis, A.-R. & McLysaght, A. Synteny-based analyses indicate that sequence divergence is not the main source of orphan genes. Elife 9, e53500 (2020).

92 Grainger, D. C. The unexpected complexity of bacterial genomes. Microbiology 162, 1167–1172 (2016).

93 Kirchberger, P. C., Schmidt, M. L. & Ochman, H. The Ingenuity of Bacterial Genomes. Annual Review of Microbiology 74, 815–834 (2020).

94 Woolstenhulme, C. J., Guydosh, N. R., Green, R. & Buskirk, A. R. High-precision analysis of translational pausing by ribosome profiling in bacteria lacking EFP. Cell Rep 11, 13–21, doi:10.1016/j.celrep.2015.03.014 (2015).

95 Livak, K. J. & Schmittgen, T. D. Analysis of Relative Gene Expression Data Using Real-Time Quantitative PCR and the 2−ΔΔCT Method. Methods 25, 402–408, doi:https://doi.org/10.1006/meth.2001.1262 (2001).

96 Hücker, S. M. et al. Discovery of numerous novel small genes in the intergenic regions of the *Escherichia coli* O157:H7 Sakai genome. PLoS One 12, e0184119, doi:10.1371/journal.pone.0184119 (2017).

97 Doellinger, J., Schneider, A., Hoeller, M. & Lasch, P. Sample Preparation by Easy Extraction and Digestion (SPEED) - A Universal, Rapid, and Detergent-free Protocol for Proteomics Based on Acid Extraction. Molecular & Cellular Proteomics 19, 209, doi:10.1074/mcp.TIR119.001616 (2020).

98 Tyanova, S., Temu, T. & Cox, J. The MaxQuant computational platform for mass spectrometry-based shotgun proteomics. Nature Protocols 11, 2301–2319, doi:10.1038/nprot.2016.136 (2016).

99 Cox, J. et al. Andromeda: a peptide search engine integrated into the MaxQuant environment. J Proteome Res 10, 1794–1805, doi:10.1021/pr101065j (2011).

100 O’Leary, N. A. et al. Reference sequence (RefSeq) database at NCBI: current status, taxonomic expansion, and functional annotation. Nucleic Acids Res 44, D733–745, doi:10.1093/nar/gkv1189 (2016).

101 MacLean, B. et al. Skyline: an open source document editor for creating and analyzing targeted proteomics experiments. Bioinformatics 26, 966–968, doi:10.1093/bioinformatics/btq054 (2010).

102 Schwanhäusser, B. et al. Global quantification of mammalian gene expression control. Nature 473, 337–342, doi:10.1038/nature10098 (2011).

103 Zolg, D. P. et al. PROCAL: A Set of 40 Peptide Standards for Retention Time Indexing, Column Performance Monitoring, and Collision Energy Calibration. PROTEOMICS 17, 1700263, doi:https://doi.org/10.1002/pmic.201700263 (2017).

104 Zuker, M. Mfold web server for nucleic acid folding and hybridization prediction. Nucleic Acids Res 31, 3406–3415, doi:10.1093/nar/gkg595 (2003).

105 Andrews, S. FastQC: a quality control tool for high throughput sequence data. Available online at: http://www.bioinformatics.babraham.ac.uk/projects/fastqc. (2010).

106 Chen, S., Zhou, Y., Chen, Y. & Gu, J. fastp: an ultra-fast all-in-one FASTQ preprocessor. Bioinformatics 34, i884–i890, doi:10.1093/bioinformatics/bty560 (2018).

107 Langmead, B. & Salzberg, S. L. Fast gapped-read alignment with Bowtie 2. Nat Methods 9, 357–359, doi:10.1038/nmeth.1923 (2012).

108 Li, H. et al. The Sequence Alignment/Map format and SAMtools. Bioinformatics 25, 2078–2079, doi:10.1093/bioinformatics/btp352 (2009).

109 Quinlan, A. R. & Hall, I. M. BEDTools: a flexible suite of utilities for comparing genomic features. Bioinformatics 26, 841–842, doi:10.1093/bioinformatics/btq033 (2010).

110 Ramírez, F. et al. High-resolution TADs reveal DNA sequences underlying genome organization in flies. Nature Communications 9, 189, doi:10.1038/s41467-017-02525-w (2018).

111 Neuhaus, K. et al. Differentiation of ncRNAs from small mRNAs in *Escherichia coli* O157:H7 EDL933 (EHEC) by combined RNAseq and RIBOseq - ryhB encodes the regulatory RNA RyhB and a peptide, RyhP. BMC genomics 18, 216–216, doi:10.1186/s12864-017-3586-9 (2017).

112 Robinson, M. D., McCarthy, D. J. & Smyth, G. K. edgeR: a Bioconductor package for differential expression analysis of digital gene expression data. Bioinformatics (Oxford, England) 26, 139–140, doi:10.1093/bioinformatics/btp616 (2010).

113 Altschul, S. F., Gish, W., Miller, W., Myers, E. W. & Lipman, D. J. Basic local alignment search tool. J Mol Biol 215, 403–410, doi:10.1016/s0022-2836(05)80360-2 (1990).

114 Kans, J. in Entrez Programming Utilities Help [Internet] (National Center for Biotechnology Information (US), 2020).

115 Buchfink, B., Xie, C. & Huson, D. H. Fast and sensitive protein alignment using DIAMOND. Nature Methods 12, 59, doi:10.1038/nmeth.3176 https://www.nature.com/articles/nmeth.3176#supplementary-information (2014).

116 Gudyś, A. & Deorowicz, S. QuickProbs 2: towards rapid construction of high-quality alignments of large protein families. Scientific reports 7, 1–12 (2017).

117 Suyama, M., Torrents, D. & Bork, P. PAL2NAL: robust conversion of protein sequence alignments into the corresponding codon alignments. Nucleic acids research 34, W609–W612 (2006).

118 Nguyen, L.-T., Schmidt, H. A., Von Haeseler, A. & Minh, B. Q. IQ-TREE: a fast and effective stochastic algorithm for estimating maximum-likelihood phylogenies. Molecular biology and evolution 32, 268–274 (2015).

119 Minh, B. Q. et al. IQ-TREE 2: New models and efficient methods for phylogenetic inference in the genomic era. Molecular Biology and Evolution 37, 1530–1534 (2020).

120 Menardo, F. et al. Treemmer: a tool to reduce large phylogenetic datasets with minimal loss of diversity. BMC bioinformatics 19, 1–8 (2018).

121 Junier, T. & Zdobnov, E. M. The Newick utilities: high-throughput phylogenetic tree processing in the UNIX shell. Bioinformatics 26, 1669–1670 (2010).

122 Huerta-Cepas, J., Serra, F. & Bork, P. ETE 3: reconstruction, analysis, and visualization of phylogenomic data. Molecular biology and evolution 33, 1635–1638 (2016).

123 Price, M. N., Dehal, P. S. & Arkin, A. P. FastTree 2–approximately maximum-likelihood trees for large alignments. PloS one 5, e9490 (2010).

124 Perez-Riverol, Y. et al. The PRIDE database and related tools and resources in 2019: improving support for quantification data. Nucleic Acids Res 47, D442–d450, doi:10.1093/nar/gky1106 (2019).

125 Sharma, V. et al. Panorama Public: A Public Repository for Quantitative Data Sets Processed in Skyline. Mol Cell Proteomics 17, 1239–1244, doi:10.1074/mcp.RA117.000543 (2018).

