## Supplementary figures for "Shadow ORFs illuminated: long overlapping genes in *Pseudomonas aeruginosa* are translated and under purifying selection"

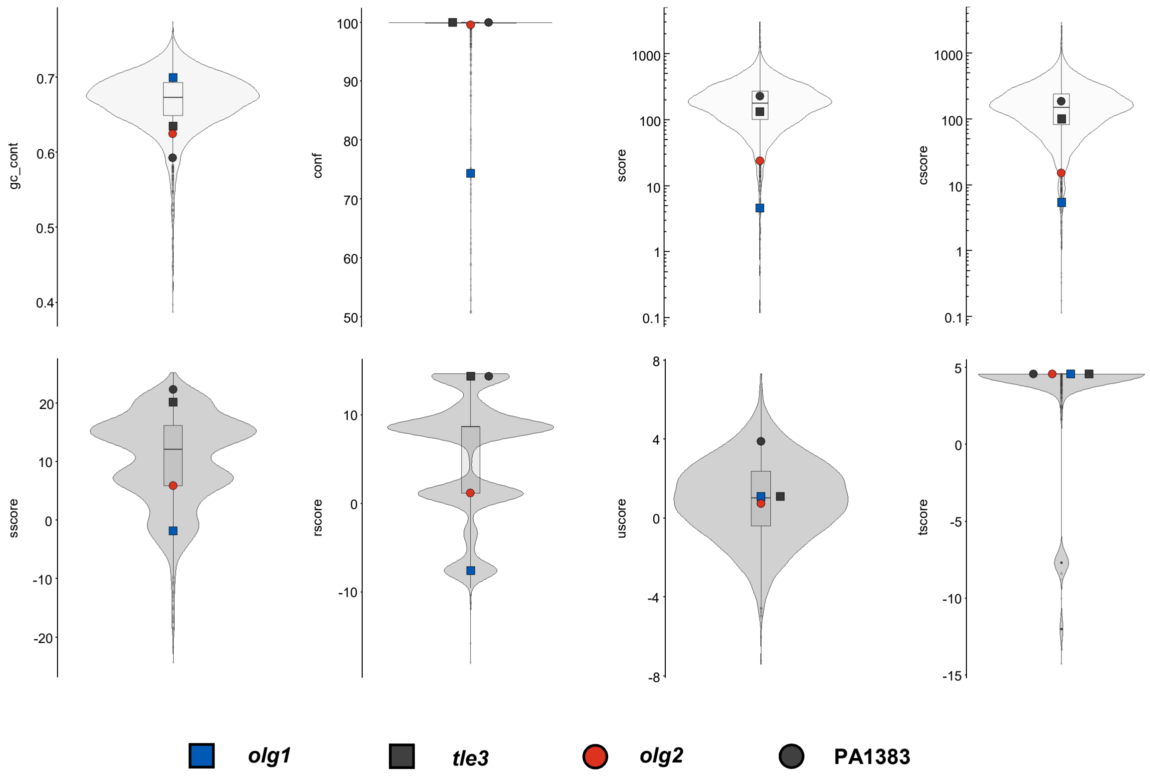


**Supplementary Fig. 1. Predicted scores of *olg1* and *olg2* in relation to all protein coding genes obtained by Prodigal (Hyatt et al.[62]).** Shown are violin plots displaying the GC content of the gene sequence (gc_cont), the confidence score (conf) indicating the likelihood of the gene to be real, the overall score (score), the hexamer coding proportion score (cscore), the translation initiation site score (sscore), the ribosome binding site score (rscore), the score for the region adjacent to the start codon (uscore) and the start codon sequence score (tscore). Included boxplots indicate 25%, 50% and 75% quartile values for all predicted, protein-coding genes (n=5,681). Values of the overlapping ORFs are represented by coloured symbols; their mother genes by the respective grey-shaded symbol.

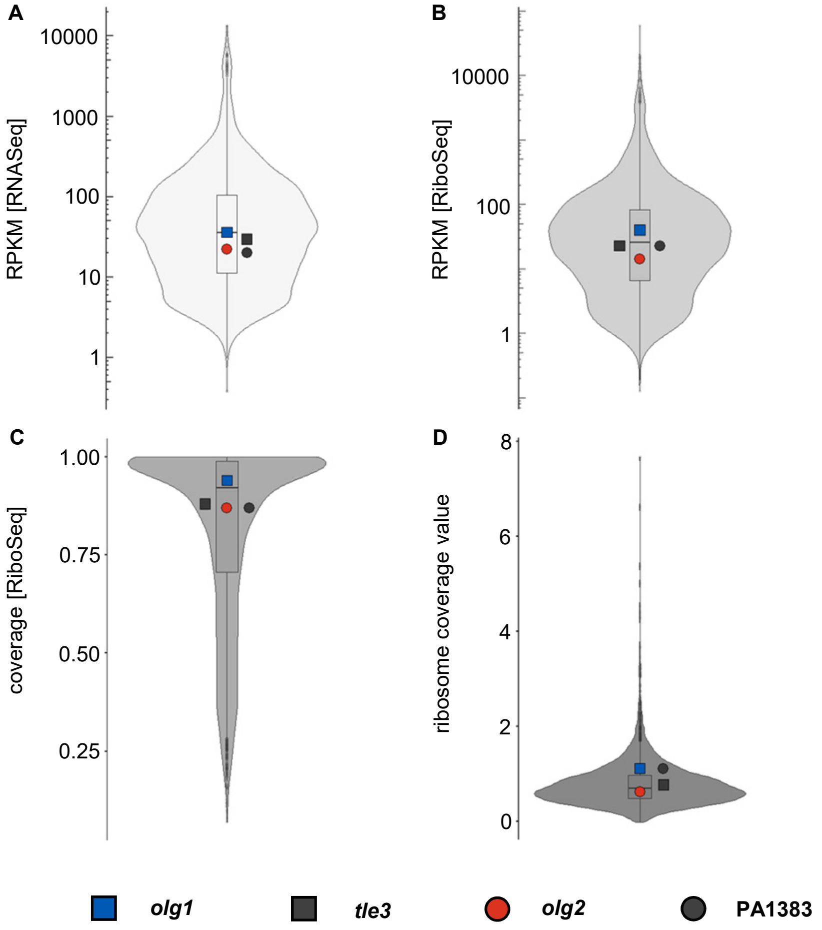


**Supplementary Fig. 2. RNASeq and RiboSeq metrics of the overlapping gene pairs *olg1*-*tle3* and *olg2*-PA1383 compared to all annotated, protein-coding genes (n=5,572).** Shown are violin plots displaying mean reads per kilobase per million mapped reads (RPKM) values for RNASeq **(A)** and RiboSeq **(B)**, mean coverage values for RiboSeq **(C)** and mean ribosome coverage values **(D)**, which are calculated by dividing the RPKM [RiboSeq] by the RPKM [RNASeq] of two biological replicates. Included boxplots indicate 25%, 50% and 75% quartile values for all annotated, protein-coding genes. Values of the overlapping ORFs are represented by coloured symbols; their mother genes by the respective grey-shaded symbol

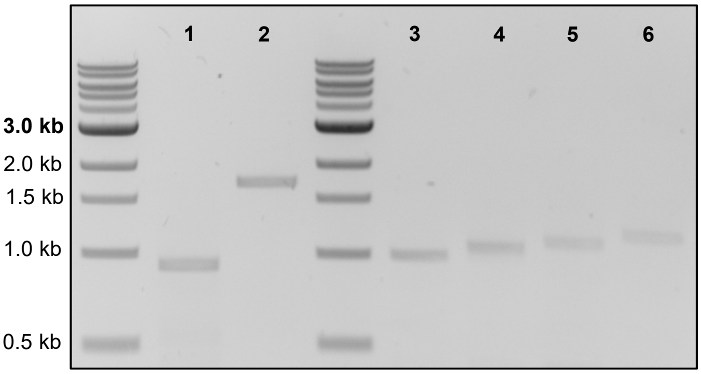


**Supplementary Fig. 3. Transcriptional signals of *olg1* and *olg2*.**
RT-PCR using primers binding at the beginning and at the end of the coding regions of *olg1* (lane 1, target length=917 nt, primer 8 & 9) and *olg2* (lane 2, target length=1696 nt, primer 10 & 11) confirmed transcription throughout the entire ORF length. Primer binding 45 nt (lane 3, target length=995 nt, Primer 15), 110 nt (lane 4, target length=1060 nt, Primer 16), 161 nt (lane 5, target length=1111 nt, Primer 17) and 240 nt (lane 6, target length=1190 nt, Primer 18) upstream of the start codon of *olg1* verified a minimum transcript length of 1190 nt.

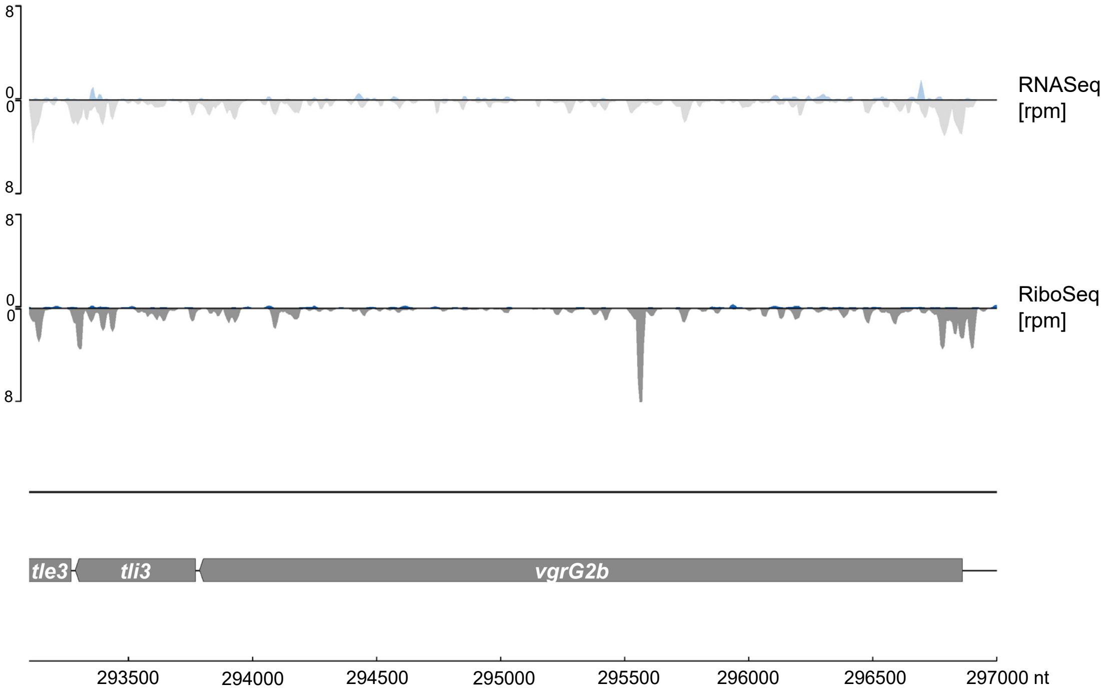


**Supplementary Fig. 4. RNASeq and RiboSeq signals upstream of the *olg1-tle3* locus.**
Shown are the mean normalized rpm values of all transcriptome (first track) and translatome reads (second track) of this study (n=2) for the annotated genes *tli3* and *vgrG2b* (both grey) located upstream of gene *tle3* (grey). Possible background reads antisense to the listed genes are highlighted in blue.

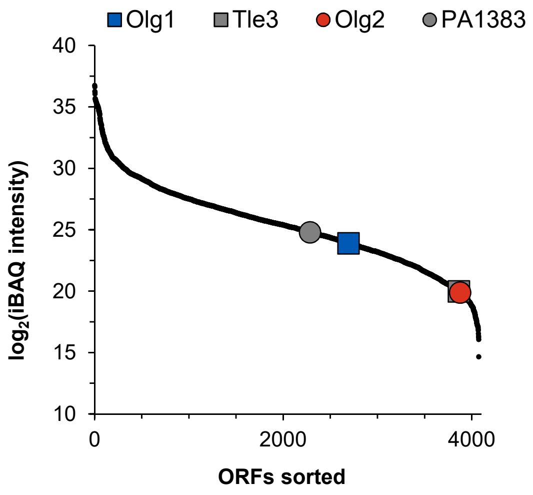


**Supplementary Figure 5. Intensities of all proteins measured by mass spectrometry in descending order.** Mass spectrometric intensities (iBAQ[102] values) of all proteins detected in the sample taken at $OD_{600nm}$=1. The proteins encoded by the OLGs are represented by coloured symbols, those of their mother genes by the respective grey-shaded symbols. Black dots represent the intensities of all quantifiable proteins encoded by annotated genes (n=4,073).

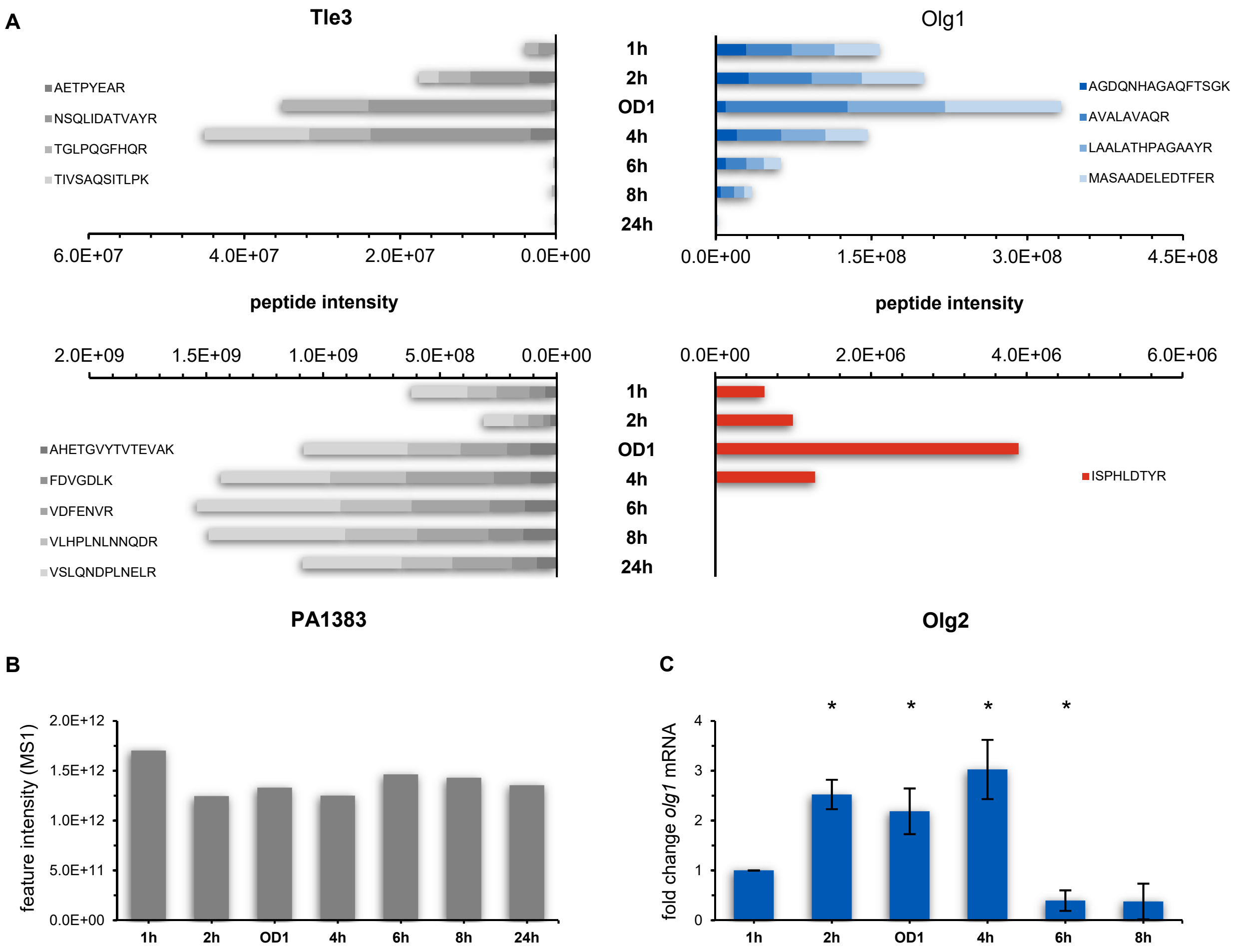


**Supplementary Fig. 6. Temporal control of Olg1 and Olg2 expression.**
**(A)** Shown are the intensities of all proteins at diverse time points during growth (1h, 2h, 4h, 6h, 8h and 24h as well as at $OD_{600nm}$=1) measured by targeted proteomics. **(B)** Loading control of the targeted proteomic experiment. Shown are the summed intensities of all measured peptides per sample. **(C)** mRNA levels measured for *olg1* via quantitative PCR. Ct values were normalized to the expression of the housekeeping gene *gyrA*. The values of the sample taken at 1 h served as a reference for fold change calculation. Shown are the mean values of three biological replicates. Comparison between the 1h sample and one of the other samples was performed by using a two-tailed Welch two sample t-test (*p ≤0.05).

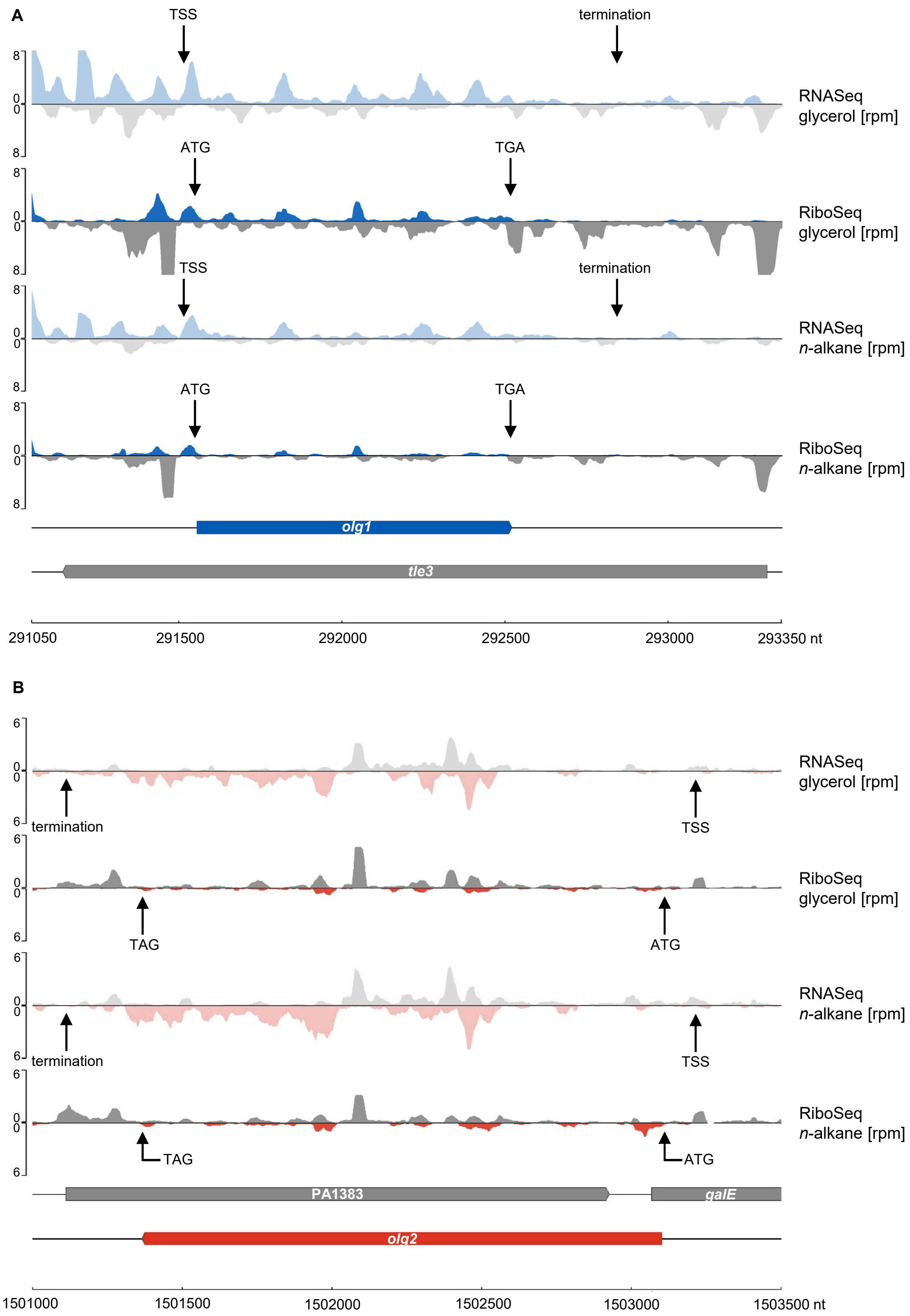


**Supplementary Fig. 7. RNASeq and RiboSeq signals at the *olg1-tle3* (A) and *olg2*-PA1383 locus (B).**
Shown are the mean normalized rpm values of all transcriptome (first & third track) and translatome reads (second & fourth track) of the datasets “M9+glycerol” and “M9+*n*-alkane” (n=3, each) published by Grady et al.[69] for *olg1* (blue), *olg2* (red) and their mother genes *tle3* and PA1383 (both grey). Transcription start (TSS) and stop sites (termination) as well as the positions of start and stop codons are indicated by arrows.

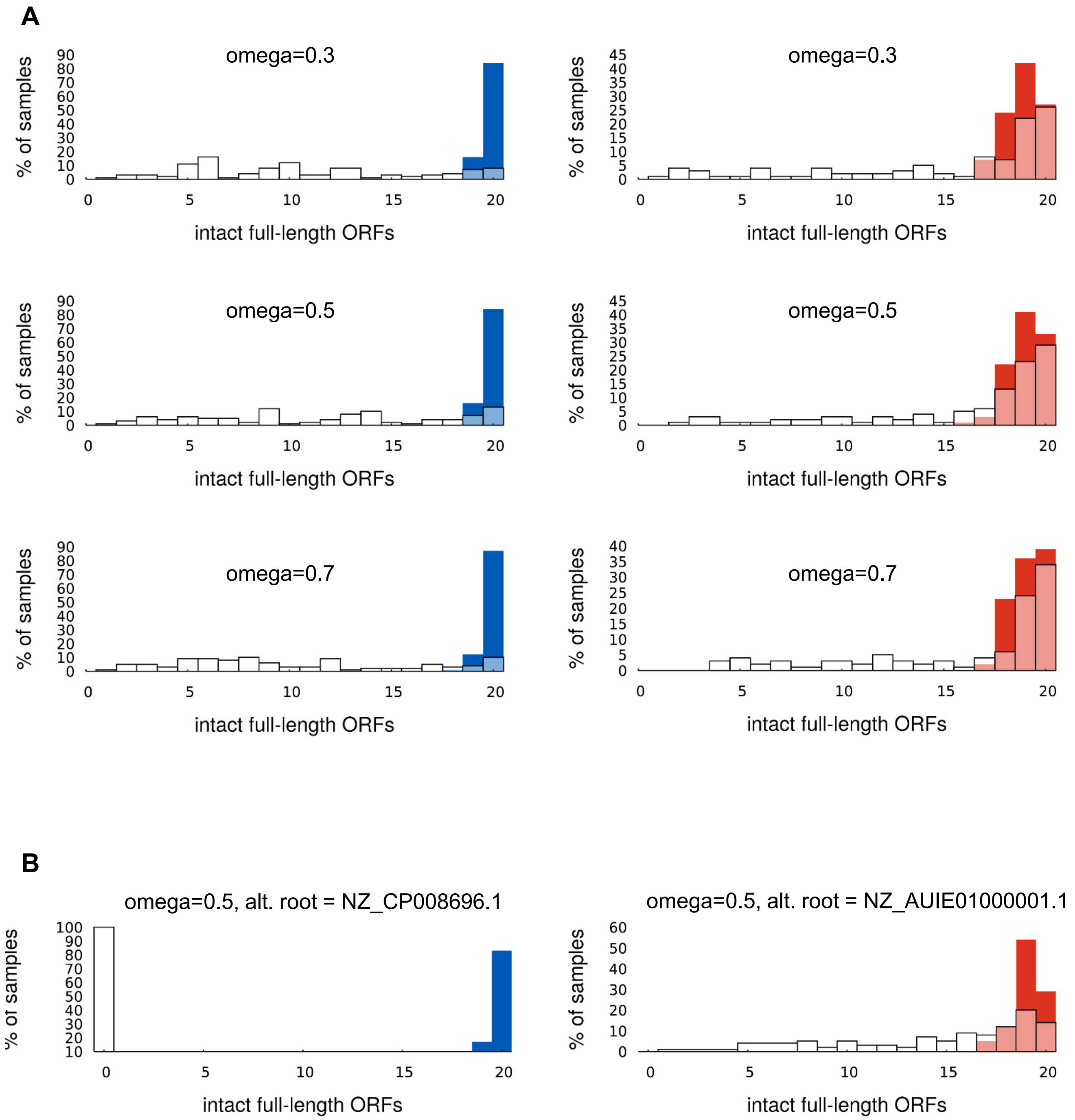


**Supplementary Figure 8. Additional evolutionary simulation analyses**
**(A)** The effect on different values of omega, which is approximately the dN/dS ratio, on the number of observed intact (no premature stop codon) ORFs in evolutionary simulations of the mother genes *tle3* (blue) and PA1383 (red). Simulations were conducted in Pyvolve. This shows only very limited effect of this parameter in these simulations. The plots for omega=0.5 are the same as in Figure 4C. **(B)** Examples of the effect of choosing a different (closer) genome as root for the tree along which evolution is simulated. It is apparent that the choice of root has a much greater effect than the variation in omega above. In these cases evidence of purifying selection against stop codons in the natural sequences is retained.
