## Supplementary information for "Shadow ORFs illuminated: long overlapping genes in *Pseudomonas aeruginosa* are translated and under purifying selection"

Five alternative start codons (CTG_291295_, TTG_291370_, CTG_291436_, ATG_291508_ & GTG_291535_) are located within the upstream region of *olg1*, providing the possibility of an N-terminal extension of the coding region of 21 to 261 nt. This hypothesis is supported by the results of RiboSeq (**Figure 2A**, second track) indicating translational signals upstream of the selected start codon. In addition, proven transcription upstream of the predicted start site (**Supplementary Figure 3**, lane 3 to 6) potentially facilitates the opportunity of a prolongated or alternative version of *olg1*. However, the following aspects argue against an N-terminal extension: Firstly, ATG is the most frequent start codon in *Pseudomonas aeruginosa*^126^ whereas the usage of alternative start codons is rather rare in prokaryotes^127^. Despite the selected start codon (ATG_291556_), only one additional ATG is located further upstream at position 291508. For this ATG, a SD and a putative promoter were predicted, but in suboptimal spacing to the start codon and with a lower probability compared to the selected ATG_291556_. With the exception of GTG_291535_, all further start codons lacked either a SD sequence or a putative promoter. However, we just tested for the presence of σ^70^ promoters and, therefore, transcription initiation driven by one of the other 23 known promoters^128^ might be conceivable. A second and probably the most descriptive aspect indicating that the coding region of *olg1* starts at the selected ATG_291556_ is provided by prediction programs. When deleting all start codons within *olg1,* Prodigal^62^ predicted ATG^291556^ to be the correct start position. Furthermore, DeepRibo^66^ also predicted translation of *olg1* starting from ATG_291556_ with the highest likelihood. Thirdly, no mass spec peptides were detected when searching against an N-terminal extended Olg1 sequence in the MS data obtained by the DDA proteomics experiment. Absence of peptides, though, does not necessarily correlate with protein presence because of various reasons, including the absence of tryptic cleavage sites^129,130^, the efficiency of protein digest and extraction^131^ or challenging peptide properties like high hydrophobicity^132^. Due to contradictory results, further experiments, e.g. frameshift mutagenesis^133^ or modified RiboSeq with translation inhibitors like tetracycline^134^ or retapamulin^81^ are necessary to unveil the correct translation start site of *olg1*.

**Supplemental references**

62	Hyatt, D. *et al.* Prodigal: prokaryotic gene recognition and translation initiation site identification. *BMC Bioinformatics* **11**, 119-119, doi:10.1186/1471-2105-11-119 (2010).

66	Clauwaert, J., Menschaert, G. & Waegeman, W. DeepRibo: a neural network for precise gene annotation of prokaryotes by combining ribosome profiling signal and binding site patterns. *Nucleic Acids Res* **47**, e36-e36, doi:10.1093/nar/gkz061 (2019).

81	Meydan, S. *et al.* Retapamulin-Assisted Ribosome Profiling Reveals the Alternative Bacterial Proteome. *Mol Cell* **74**, 481-493.e486, doi:10.1016/j.molcel.2019.02.017 (2019).

126	West, S. E. & Iglewski, B. H. Codon usage in Pseudomonas aeruginosa. *Nucleic Acids Res* **16**, 9323-9335, doi:10.1093/nar/16.19.9323 (1988).

127	Bachvarov, B., Kirilov, K. & Ivanov, I. Codon Usage in Prokaryotes. *Biotechnology & Biotechnological Equipment* **22**, 669-682, doi:10.1080/13102818.2008.10817533 (2008).

128	Potvin, E., Sanschagrin, F. & Levesque, R. C. Sigma factors in Pseudomonas aeruginosa. *FEMS Microbiology Reviews* **32**, 38-55, doi:10.1111/j.1574-6976.2007.00092.x (2008).

129	Slavoff, S. A. *et al.* Peptidomic discovery of short open reading frame–encoded peptides in human cells. *Nature Chemical Biology* **9**, 59-64, doi:10.1038/nchembio.1120 (2013).

130	Landry, C. R., Zhong, X., Nielly-Thibault, L. & Roucou, X. Found in translation: functions and evolution of a recently discovered alternative proteome. *Current Opinion in Structural Biology* **32**, 74-80, doi:https://doi.org/10.1016/j.sbi.2015.02.017 (2015).

131	Baldwin, M. A. Protein Identification by Mass Spectrometry. *Molecular &amp;amp; Cellular Proteomics* **3**, 1, doi:10.1074/mcp.R300012-MCP200 (2004).

132	Bagag, A. *et al.* Characterization of hydrophobic peptides in the presence of detergent by photoionization mass spectrometry. *PLoS One* **8**, e79033, doi:10.1371/journal.pone.0079033 (2013).

133	Smollett, K. L. *et al.* Experimental determination of translational start sites resolves uncertainties in genomic open reading frame predictions - application to Mycobacterium tuberculosis. *Microbiology (Reading, England)* **155**, 186-197, doi:10.1099/mic.0.022889-0 (2009).

134	Nakahigashi, K. *et al.* Comprehensive identification of translation start sites by tetracycline-inhibited ribosome profiling. *DNA Research* **23**, 193-201, doi:10.1093/dnares/dsw008 (2016).
